# A truncated SARS-CoV-2 nucleocapsid protein enhances virus fitness by evading antiviral responses

**DOI:** 10.1101/2025.02.15.638421

**Authors:** Rory P. Mulloy, Danyel Evseev, Maxwell P. Bui-Marinos, Noga Sharlin, Jennifer A. Corcoran

**Affiliations:** Microbiology, Immunology and Infectious Diseases Department, Snyder Institute for Chronic Disease, Charbonneau Cancer Research Institute, Cumming School of Medicine, University of Calgary, Calgary, Alberta, Canada

## Abstract

Viruses face a selective pressure to evade cellular antiviral responses to control the outcome of an infection. However, due to their limited genome size, viruses must adopt unique strategies to confront cellular sensors. Since emerging in humans, SARS-CoV-2 has accrued multiple mutations throughout its genome, some of which enhanced virus replication and led to the rise of viral variants. However, the biological consequences of many of these changes remain to be discovered. Here, we show that SARS-CoV-2 produces a truncated form of the nucleocapsid protein, called N*^M210^. Due to the acquisition of a viral transcription regulatory sequence (TRS) in the N gene, certain variants, such as Omicron, produce a new viral mRNA that markedly increases N*^M210^ expression. We show that N*^M210^ is a dsRNA binding protein, which inhibits multiple arms of the cellular antiviral response, including blocking interferon induction and inhibiting stress granule formation. We created a panel of recombinant SARS-CoV-2 viruses (rSARS-2) with mutations in the N gene that increased or decreased N*^M210^ production. We show that N*^M210^ production increases virus fitness, as viruses that produce more N*^M210^ outcompeted wild-type rSARS-2. We demonstrate that the fitness advantage provided by N*^M210^ is partly due to its ability to potently block stress granules. We propose a model where, to evade the cellular antiviral response, SARS-CoV-2 has evolved a mechanism to increase the production of a truncated form of the N protein, which broadly limits the activation of dsRNA-induced antiviral responses, tipping the balance in favour of the virus in the battle for control of the cell.

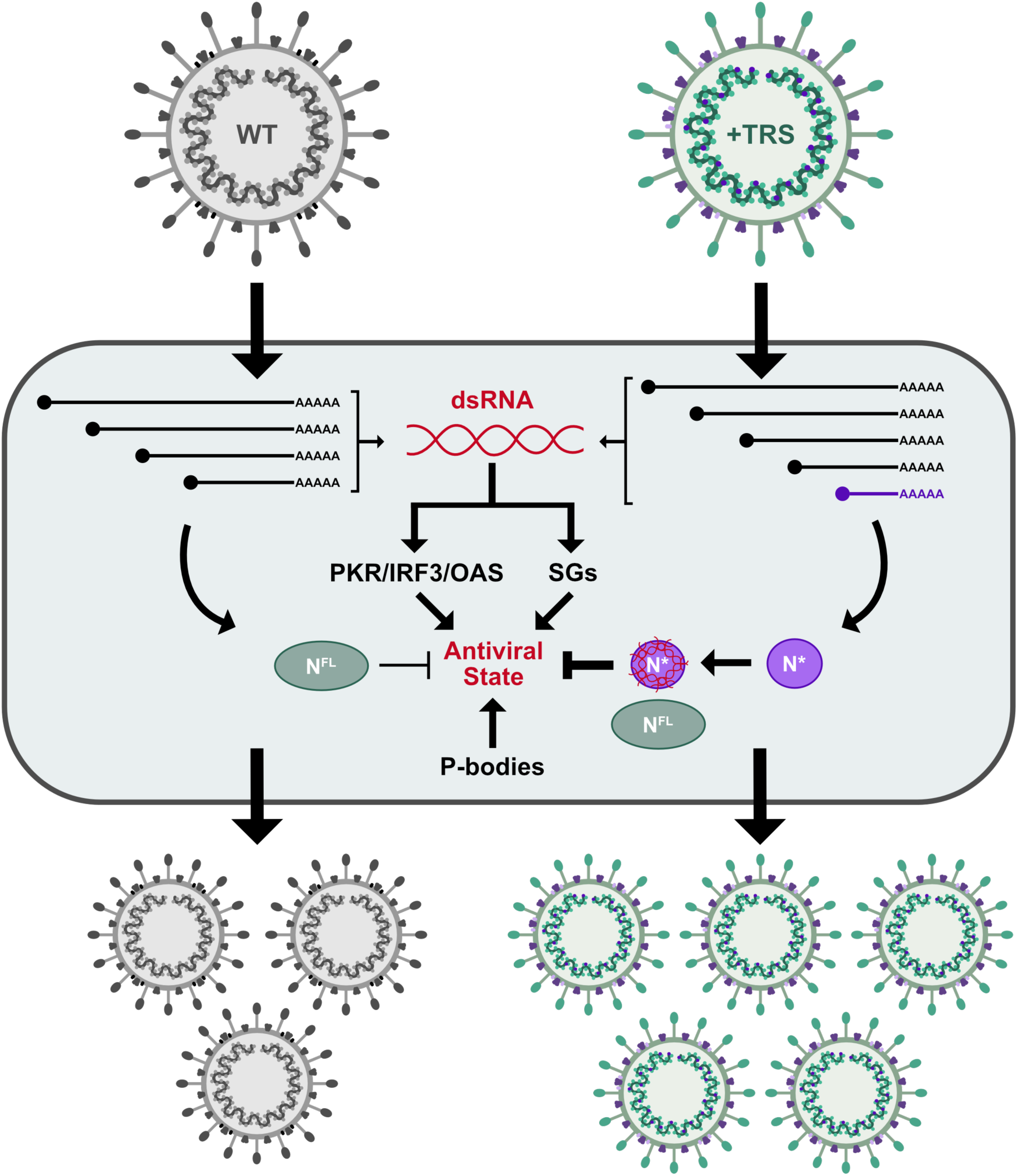

**Highlights:**

- SARS-CoV-2 variants evolved to upregulate truncated N (N*) synthesis to increase virus fitness
- N*^M210^ is a potent dsRNA-binding protein that blocks cellular dsRNA sensing
- N*^M210^ inhibits stress granule formation independent of G3BP1 binding

## INTRODUCTION

The outcome of a viral infection is determined by two factors: the ability of the cell to sense and respond to the virus, and the ability of the virus to evade these responses. Through this evolutionary conflict, cells have developed numerous distinct mechanisms to restrict viral replication, and in turn viruses have acquired methods to suppress these antiviral responses to promote their replication and spread. Many viruses, including coronaviruses (CoVs), produce double-stranded RNA (dsRNA) during infection, activating multiple antiviral responses (V’kovski et al., 2021; Weber et al., 2006). For example, dsRNA sensing induces translational arrest and the production of type I interferons (IFN), triggering a paracrine and autocrine induction of an antiviral state (Steiner et al., 2024; V’kovski et al., 2021). More recently, antiviral programmes that rely on the formation of ribonucleoprotein (RNP) granules, including stress granules (SGs) and processing bodies (P-bodies) have been described (McCormick & Khaperskyy, 2017; Reineke & Lloyd, 2013). Although this is a rapidly emerging field, the precise mechanism for how these RNP granules antagonize virus replication remains incompletely understood. What is clear is that a diversity of viruses, including CoVs, potently and consistently inhibit SGs and P-bodies during infection (Corcoran et al., 2012, 2015; Dolliver et al., 2022; Dougherty et al., 2015; Fan et al., 2021; Gaete-Argel et al., 2019; Kleer. et al., 2022; Long et al., 2024; McCormick & Khaperskyy, 2017; Rabouw et al., 2016). Both the interferon-based and ‘granular’ antiviral pathways impose a barrier to virus replication, explaining why viruses have evolved mechanisms to evade or dampen these antiviral programmes. As viruses replicate in a new host, there is a selective pressure to optimize their evasion strategies. Viruses with a superior ability to evade cellular antiviral responses will have a fitness advantage, and thus will be more likely to replicate and spread (Carabelli et al., 2023; Park & Iwasaki, 2020; Thorne et al., 2022).

Severe-acute respiratory syndrome CoV-2 (SARS-CoV-2) is an RNA virus and is the causative agent of coronavirus disease 2019 (COVID-19). Despite concerted efforts to produce vaccines and antivirals, due to its evolutionary dexterity, SARS-CoV-2 still poses a significant health risk. The pathogenicity of SARS-CoV-2 has been largely attributed to a mismanaged immune response; the virus potently represses the antiviral response activation, resulting in a delayed and exacerbated cytokine production late in infection (Hadjadj et al., 2020; Hawerkamp et al., 2023; Lucas et al., 2020; Zhang et al., 2020). Following emergence and circulation in the human population, SARS-CoV-2 accrued a multitude of mutations throughout its genome, some of which have improved virus fitness, enabling new genotypes to outcompete the ancestral virus, driving the rise of viral variants (Carabelli et al., 2023; Thorne et al., 2022). While much of the attention has been focused on changes in the spike gene, many other SARS-CoV-2 proteins contribute to viral transmission, pathology, and overall virus fitness; however, the biological consequences of most of these changes remain undiscovered. One viral gene subject to significant evolutionary change is the nucleocapsid (N) gene (Johnson et al., 2022; Kubinski et al., 2024; Syed et al., 2021; H. Wu et al., 2021). Here, we focus on one genetic change in the N gene and determine how its acquisition impacts SARS-CoV-2 replication, fitness, and antiviral antagonism.

Upon infection, SARS-CoV-2 delivers its 30 kb genome to the cytoplasm where it is directly translated, promoting the synthesis of non-structural proteins (nsps), including the viral RNA polymerase and proteins that form replication organelles (Steiner et al., 2024). In replication organelles, SARS-CoV-2 transcribes viral messenger RNAs called subgenomic RNAs (sgRNAs) (Steiner et al., 2024). sgRNA synthesis is governed by the presence of short ∼6 nucleotide sequences, called transcription regulatory sequences (TRSs; black boxes in Fig 1A) in the genomic RNA (Sawicki & Sawicki, 1998; Sola et al., 2011; Van Marle et al., 1999). The leader TRS (L-TRS) is located at the 5’ end of the genome, whereas the body TRS (B-TRS) sequences are positioned near the 5’ end of some accessory genes and each structural gene, including N (Fig 1A). Certain SARS-CoV-2 variants have acquired a TRS in the gene body of N, resulting in the synthesis of a new sgRNA, which is thought to code for an amino-terminally truncated form of the N protein (Mears et al., 2025; Thorne et al., 2022); however, the precise function of this truncated product remains unclear.

**Figure 1.**
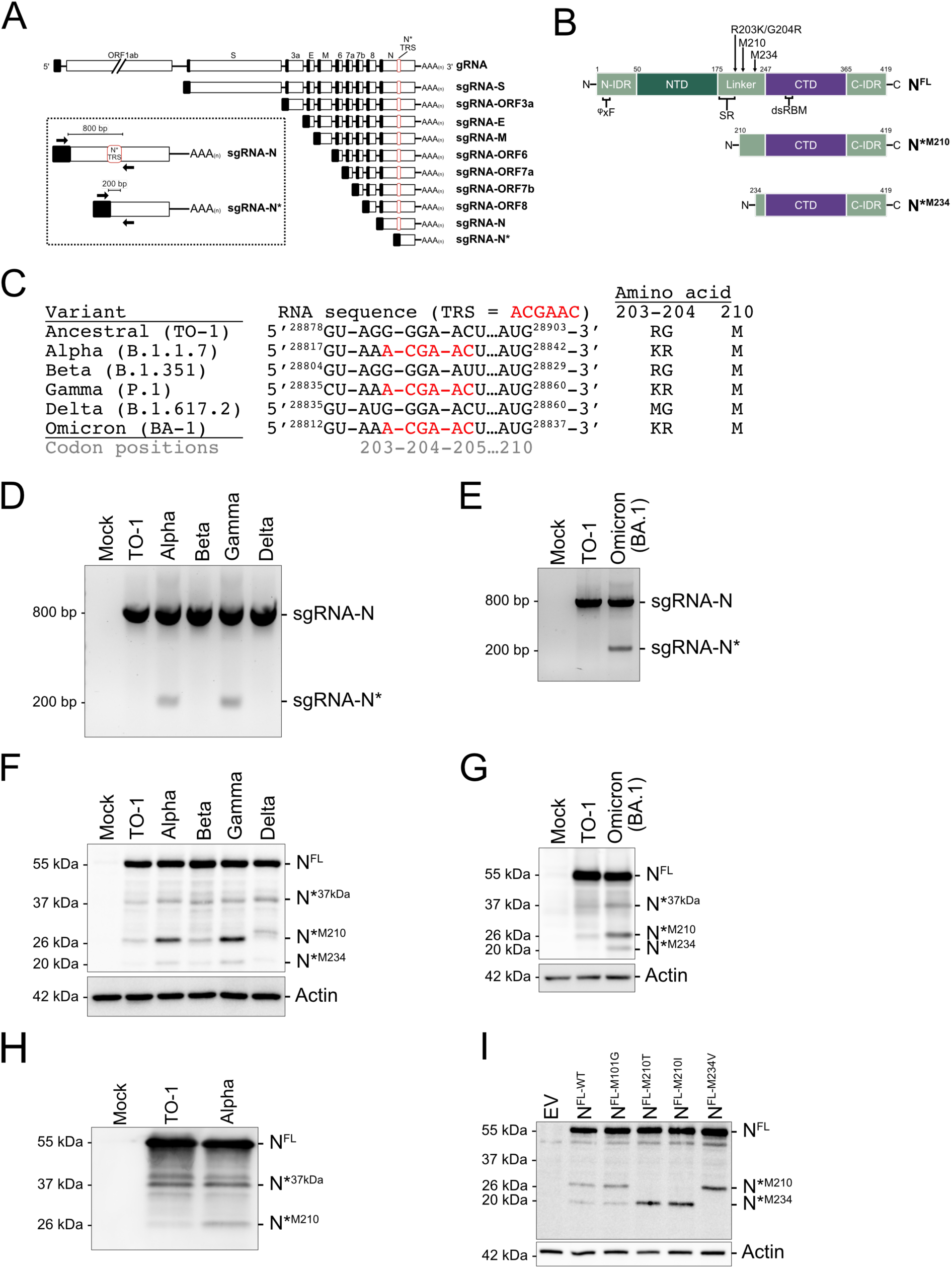
Select SARS-CoV-2 variants produce truncated proteoforms of the nucleocapsid protein by internal TRS acquisition. **A.** A schematic representation of the genomic and subgenomic RNAs with canonical TRSs (black boxes) and the novel TRS (red box). The inset depicts the strategy to detect sgRNA-N* by RT-PCR. Forward primer binds to the 5’end of the leader sequence (40-75 nt) and the reverse primer binds down-stream of the novel TRS (29035-29053 nt). **B.** Domain organization of N^FL^, and truncated N proteoforms N*^M210^ and N*^M234^. Amino acid positions are labeled above, relative to nomenclature for N^FL^. Intrinsically disordered region (IDR), N-terminal domain (NTD), and C-terminal domain (CTD), serine-arginine rich area (SR), double-stranded RNA binding motif (dsRBM). φxF (φ is a hydrophobic amino acid; I15-F17) is the G3BP1-binding motif (Biswal, Lu, and Song 2022). **C.** SARS-CoV-2 variant sequence diversity at the mutational hotspot in the N gene where canonical TRS insertion occurred. Sequence of the TRS hexamer is labeled in red. Codon positions based on the ancestral sequence. **D-E.** sgRNA profile of SARS-CoV-2 variants was determined by infecting Calu3 cells (MOI=2 for D, MOI=1 for E) and RNA was harvested 24 hpi and subjected to RT-PCR and agarose gel electrophoresis. Toronto-1 (TO-1) isolate is the Wuhan-like ancestral variant. **F-G.** N proteoform profile of SARS-CoV-2 variants was determined by infecting Calu3 cells (MOI=2 for F, MOI=1 for G). Protein lysate was harvested 24 hpi and subjected to SDS-PAGE and immunoblotting with anti-N and anti-actin antibodies. **H.** N proteoform profile of extracellular SARS-CoV-2 virions. VeroE6 cells were infected with TO-1 or Alpha (MOI=1). Supernatant was harvested 24 hpi and purified via ultracentrifugation. Concentrated virions were lysed and resolved by SDS-PAGE and immunoblotting. **I.** HEK293Ts were transfected with plasmids encoding a carboxy-terminally FLAG-tagged versions of N^FL^ either without substitution of internal methionine codons (N^FL-WT^) or with the indicated internal methionine codon substitutions or with an empty vector (EV) control. Protein lysates were harvested 24 hours post-transfection and resolved by SDS-PAGE and immunoblotting for FLAG and actin.

The N protein is a remarkably multifunctional protein with essential roles in CoV replication, such as transcription, genome replication and packaging (McBride et al., 2014; W. Wu et al., 2023). Additionally, N blocks multiple branches of the cellular antiviral response, preventing translation arrest, inhibiting interferon production, and blocking SGs and P-bodies (Dolliver et al., 2022; Kleer. et al., 2022; LeBlanc et al., 2023; Oh & Shin, 2021; Zheng et al., 2021). Structurally, N is a modular protein containing two independently folded domains: the N-terminal domain (NTD) and the C-terminal domain (CTD). These domains are flanked and linked by flexible disordered regions (N-IDR, linker, and C-IDR) (Fig 1B). Both the NTD and the CTD contain RNA-binding activity, each with unique specificities; the NTD binds TRS-like sequences with sequence specificity, whereas the CTD has a high affinity for structured and dsRNA without known sequence specificity (Iserman et al., 2020; Roden et al., 2022).

Here, we show that SARS-CoV-2 produces multiple truncated versions of N (N*). One truncated N proteoform is made by internal translation initiation at the methionine codon at position 210 of the full-length N sequence and is referred to as N*^M210^. Moreover, multiple viral variants, including Omicron, upregulate N*^M210^ production due to the acquisition of a canonical TRS within the N gene. This internal TRS permits the transcription of a novel sgRNA, from which N*^M210^ is efficiently translated. Using a panel of recombinant SARS-CoV-2 viruses, we reveal that enhanced N*^M210^ production increases virus fitness. We show that N*^M210^ sequesters dsRNA and prevents activation of dsRNA-induced antiviral responses. Furthermore, N*^M210^ blocks antiviral RNPs, SGs and P-bodies by interacting with structured or dsRNAs. Using competition assays, we demonstrate that the fitness advantage provided by N*^M210^ production is, in part, due to its potent ability to block SG formation. By characterizing the function of this novel viral gene product, we explain how a genetic adaptation likely contributed to the dominance of certain variants, giving these viruses the edge to overcome the host response and spread.

## RESULTS

### SARS-CoV-2 produces truncated N proteoforms

The nucleocapsid (N) gene has been under considerable evolutionary pressure, with unique mutations arising in all the dominant variants, including Alpha (B.1.1.7), Beta (B.1.351), Gamma (P.1), Delta (B.1.617.2), and Omicron (BA-1). The linker region in N contains a mutational hotspot, resulting in various amino acid changes compared to the ancestral virus; including, R203M in the Delta lineage and R203K/G204R in Alpha, Gamma, and Omicron lineages (Fig 1B & C). We and others noticed that codon change producing the R203K/G204R substitution forms a canonical transcription regulatory sequence (TRS) within the gene body of N (Fig 1C) (Mears et al., 2025; Thorne et al., 2022). To test if this internal TRS acquisition enables transcription of a novel sgRNA, we isolated RNA from SARS-CoV-2-infected cells and conducted RT-PCR diagnostics. Using a forward primer complementary to the L-TRS (found at the 5’ end of all sgRNAs) and a reverse primer complementary to sequences in the body of the N gene, PCR amplification will produce a ∼800 bp amplicon when hybridized to canonical full-length N sgRNA and a ∼200 bp amplicon if hybridized to the novel truncated sgRNA (Fig 1A inset). We could detect amplicons corresponding to the canonical full-length sgRNA-N from all SARS-CoV-2 variants and the Wuhan-like ancestral virus, TO-1 (Fig 1D & E). However, only variants that contain the internal TRS produced the ∼200 bp amplicon expected for the novel sgRNA. These data confirm that the novel TRS promotes the transcription of a new sgRNA, sgRNA-N*.

To determine if sgRNA-N* is protein-coding, we looked for open-reading frames. The N gene contains several in-frame methionine-coding codons (AUGs) with the potential to produce truncated N proteoforms by internal translation initiation. These methionine (M) codons can be found at position 101, 210, and 234. M210 and M234 are located immediately downstream of the internal TRS, which denotes the transcription start site for sgRNA-N*, suggesting that they could be efficiently translated from this novel sgRNA (Fig 1B). To determine if SARS-CoV-2 infection produces truncated N proteoforms, we infected cells with the panel of viral variants and conducted immunoblotting for N. We found that all viruses tested produced multiple lower molecular weight proteoforms of N, including 37 kDa, 26 kDa, and 20 kDa products (Fig 1F & G). Two pieces of data indicate that the 26 kDa band corresponds to an amino-terminally truncated N proteoform initiating at M210, which will hereafter be referred to as N*^M210^: first, over-expression of a truncated N proteoform starting at M210 is 26 kDa (Fig S1A); second, this product was upregulated by variants that produce sgRNA-N* (Fig 1D & E). Variants that do not produce sgRNA-N* (TO-1, Beta, and Delta), still produce N*^M210^ protein, albeit to a much lower amount (Fig 1F & G). This is likely a result of internal ribosomal initiation at the downstream AUG at M210. Given that N^FL^ is a structural protein, we wondered if N*^M210^ could also be packaged into viral particles. To test this, extracellular virions from TO-1 and Alpha-infected cells were concentrated, lysed, and subjected to immunoblotting. We determined that multiple N* proteoforms, including N*^M210^, can be found in viral particles (Fig 1H).

We also observed a 20 kDa proteoform was produced by all variants tested but upregulated in cells infected with internal TRS-containing viruses (Fig 1F & G). This product likely derives from internal translation initiation at the AUG codon in position M234 (Fig S1A & B); hereafter this N proteoform will be referred to as N*^M234^. N*^M234^ is minimally produced by internal ribosomal initiation from the sgRNA-N transcript; however, viruses with the internal TRS (Alpha, Gamma, Omicron) have increased N*^M234^ expression via internal initiation from the shorter sgRNA-N* transcript. When N^FL^ was ectopically expressed in HEK293T cells, we could detect a similar array of lower molecular weight proteoforms, including N*^M210^ and N*^M234^, supporting the hypothesis that N* products can be produced by internal translation initiation (Fig 1I). The 37 kDa product (termed N*^37kDa^) was not detected during ectopic expression. Furthermore, substitution of M210 and M234 in the N^FL^ sequence caused the disappearance of N*^M210^ and N*^M234^, respectively (Fig 1I). Substitution of M101 had no effect on the production of truncated N proteoforms. These data suggest that N*^M210^ and N*^M234^, but not N*^37kDa^ are produced by internal ribosomal initiation. To uncouple the production of N^FL^ from N* during ectopic expression experiments, we mutated downstream initiation codons in N^FL^ (N^FL-M210I/M234V^) and N*^M210^ (N*^M210-M234V^) for all subsequent experiments unless otherwise noted (Fig S1A & B). To determine the kinetics of sgRNA-N* transcription and N* protein production, Calu3 cells were infected with the Alpha variant and RNA and protein lysates were harvested at various times post-infection. We observed that sgRNA-N and sgRNA-N* were produced concurrently, with transcription initiating between 2-4 hours post-infection (hpi). N^FL^ and N*^M210^ proteins were both first detected between 4-8 hpi (Fig S1C). Taken together, these data show that SARS-CoV-2 produces truncated proteoforms (N*) in two distinct ways: i) by internal translation initiation from in-frame AUGs; and ii) by the synthesis of a novel sgRNA in select variants. Given that SARS-CoV-2 appears to have acquired the ability to produce increased amounts of truncated N proteoforms, especially N*^M210^, this begged the question, does N* production increase virus fitness?

### Increased N*^M210^ production confers a viral fitness advantage

Prior studies reported that the R203K/G204R substitution enhanced viral fitness by increasing N^FL^ phosphorylation (Johnson et al., 2022; H. Wu et al., 2021). However, this mutation also promotes the synthesis of sgRNA-N*, which increases the production of N*^M210^ (Fig 1). Using the ancestral wild-type (WT) backbone (Chiem et al., 2021; C. Ye et al., 2020), we constructed a panel of recombinant SARS-CoV-2 viruses (rSARS2) designed to uncouple the effects of the R203K/G204R amino acid change from the insertion of the novel TRS. First, we created a virus containing the R203K/G204R substitution with the authentic nucleotide sequence, resulting in the insertion of the canonical TRS; this virus was named KR^+TRS^ (Fig 2A). Next, we created a virus containing the R203K/G204R substitution without the introduction of a TRS utilizing the degeneracy of the amino acid code; this virus was named KR^-TRS^. To block all N*^M210^ production via internal ribosome initiation, we also substituted the N*^M210^ start codon in the WT backbone; this virus was named M210I. Each recombinant virus was validated by whole genome sequencing. With this rSARS2 panel, we aimed to test if N*^M210^ synthesis could influence viral fitness.

**Figure 2.**
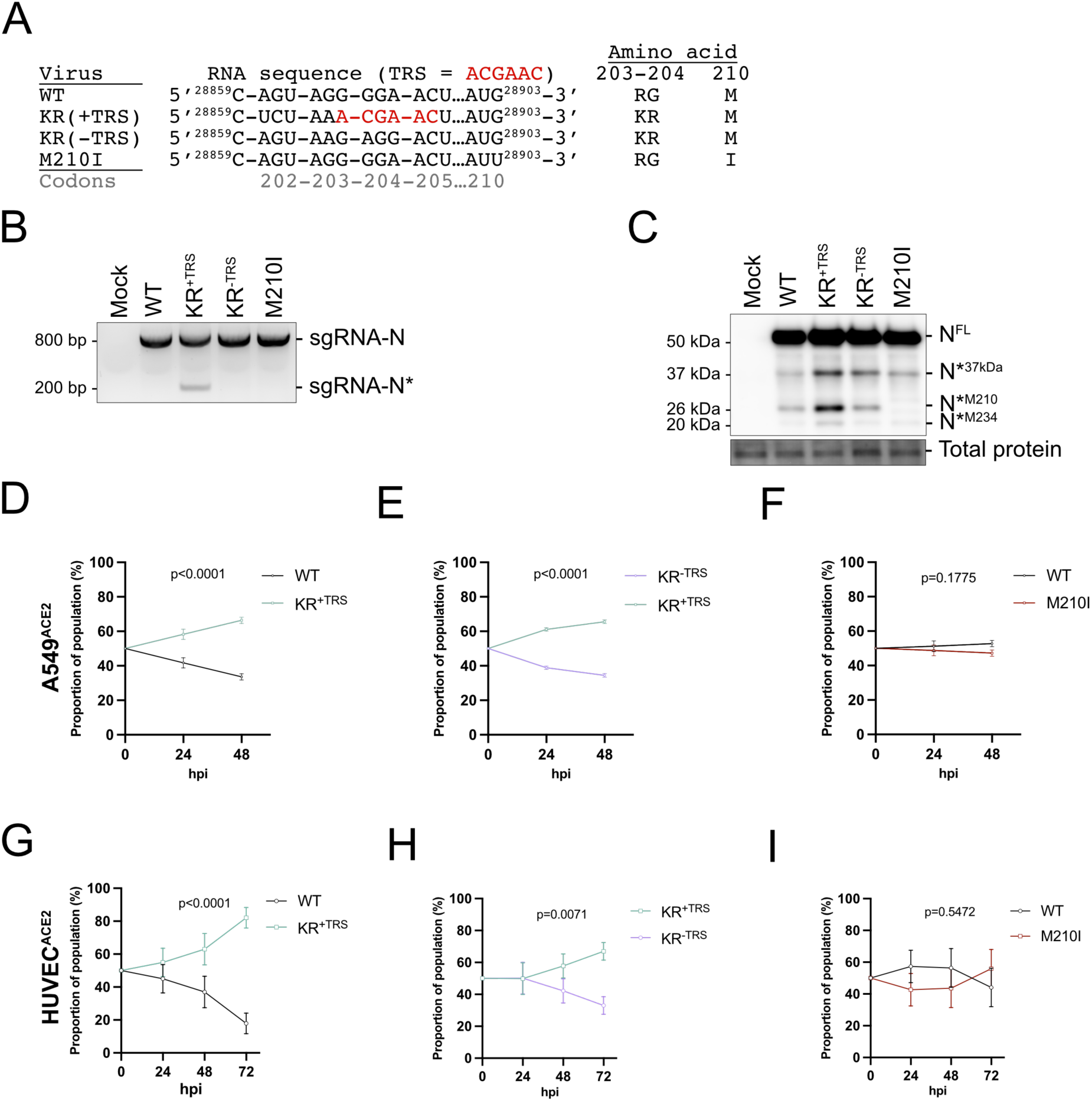
Production of N*^M210^ increases recombinant SARS-CoV-2 fitness. **A.** Sequences of recombinant SARS-CoV-2 in the mutational hotspot in the N gene. Canonical TRS hexamer labeled in red. The sequences flanking the TRS in KR^+TRS^ were based on the Gamma variant. **B.** sgRNA profile of recombinant SARS-CoV-2 was determined by infecting A549^ACE2^ cells (MOI=4). RNA was harvested 24 hpi and subjected to RT-PCR and agarose gel electrophoresis. **C.** N proteoform profile of SARS-CoV-2 variants was determined by infecting A549^ACE2^ cells (MOI=4). Protein lysate was harvested 24 hpi and subjected to SDS-PAGE and immunoblotting with anti-N. **D-I.** A549^ACE2^ cells (E-G) or primary HUVEC^ACE2^ cells (H-J) were coinfected with equal infectious titers of the indicated recombinant virus to achieve a total MOI of 0.1 for A549^ACE2^ or MOI of 0.02 for HUVEC^ACE2^. Time 0 represents the inferred proportions of each recombinant based on infectious titer input. Intracellular RNA was harvested at the indicated times post-infection and subjected to probe-based RT-qPCR to differentiate recombinant virus abundance. n=3 (E-F), n=6 (G-I). A simple linear regression was conducted, standard error mean; SEM.

We used diagnostic RT-PCR to validate that insertion of the TRS in the KR^+TRS^ virus promoted sgRNA-N* synthesis. Indeed, we found that only KR^+TRS^ produced the 200 bp amplicon indicative of sgRNA-N*, whereas all recombinants produced the 800 bp amplicon, corresponding to sgRNA-N (Fig 2B, S2A). We also observed that cells infected with KR^+TRS^ produced N*^M210^ and N*^M234^ in greater abundance than WT and KR^-TRS^ recombinants, while M210I did not produce any N*^M210^ (Fig 2C, S2B). The inability to detect the N*^M210^ proteoform from M210I-infected cells strongly suggests that N*^M210^ is produced via internal ribosomal initiation and not by proteolytic cleavage, as previously suggested (Chu et al., 2022). We observed that the 37 kDa proteoform (N*^37kDa^) was produced in cells infected with all recombinants, though it was more abundant in cells infected with KR^+TRS^ and KR^-TRS^. The precise identity of N*^37kDa^ remains unknown.

To test if increased production of N*^M210^ provides a fitness advantage, we performed competition assays between recombinant viruses that make more N*^M210^ (KR^+TRS^) compared to viruses that make less or no N*^M210^ (WT, KR^-TRS^, and M210I). Recombinant virus species were added in a 1:1 ratio based on infectious titer to A549^ACE2^ cells and primary endothelial cells (HUVEC^ACE2^). Intracellular RNA was harvested at various times post-infection. The abundance of each recombinant virus species was quantified using a probe-based RT-qPCR assay where the probe binding site spans the mutated region of the TRS or M210I substitution, thereby allowing each recombinant virus to be differentiated in a mixed population and the relative proportion of each viral RNA species to be determined (Fig S2C). Probes were validated for specificity with intracellular RNA extracted from cells independently infected with each recombinant virus (Fig S2D).

In both cell types, KR^+TRS^ readily outcompeted the WT virus. In A549^ACE2^ cells, 66 percent of the virus reads corresponded to KR^+TRS^ viral RNA by 48 hpi, and in HUVEC^ACE2^ cells, 82 percent of the virus reads corresponded to KR^+TRS^ by 72 hpi (Fig 2E & H). To discern whether the fitness advantage derived from the amino acid change R203K/G204R or the internal TRS nucleotide acquisition, which are both present in the KR^+TRS^ virus, we next conducted competition assays between KR^+TRS^ and KR^-TRS^ recombinant viruses. These viruses produce identical N^FL^ proteins but differ in the abundance of N*^M210^ that is made during infection (Fig 2C). In both A549^ACE2^ and HUVEC^ACE2^ cells, the KR^+TRS^ virus outcompeted the KR^-TRS^ virus, suggesting that greater N*^M210^ production confers a fitness advantage (Fig 2E & H). Given that N*^M210^ can be basally produced by internal ribosome initiation at M210 by both WT and KR^-TRS^ viruses, we wondered if even low levels of N*^M210^ could provide a fitness advantage. To test this, we performed competition assays between WT and M210I viruses. However, we detected no significant fitness differences between these viruses, suggesting that basal production of N*^M210^ does not significantly alter viral fitness *in vitro* (Fig 2F & I). During co-infection competition assays, we found that KR^+TRS^ outcompeted WT and KR^-TRS^ from intracellular RNA analysis; therefore, we expected that this fitness advantage would result in more KR^+TRS^ virions being produced and secreted into the extracellular space. We found that extracellular RNA ratios were similar to those seen with intracellular RNA analysis, as the KR^+TRS^ virus outcompeted both WT and KR^-TRS^ viruses, whereas WT and M210I were not statistically different (Fig S2E-G). Taken together, these data show that viruses that produce N*^M210^ in greater abundance have enhanced fitness, yet the mechanism by which N*^M210^ promotes virus replication is unknown.

### N*^M210^ is a dsRNA-binding protein

The production of a truncated version(s) of a parent protein can significantly alter protein function by changing localization, post-translational modifications, or interactors. We speculated that loss of the amino terminal 209 amino acids of N^FL^ licensed N*^M210^ with unique functions that support viral replication and/or immune evasion. N^FL^ was shown to possess double-stranded RNA (dsRNA) binding activity, thought to be necessary to support viral RNA synthesis, packaging, and antagonism of the antiviral response, all features that would promote efficient virus infection (Aloise et al., 2023; Iserman et al., 2020; Li et al., 2021; Roden et al., 2022). dsRNA-binding activity is dependent on two lysine residues, K257 and K261, in the dsRNA binding motif (dsRBM) (Aloise et al., 2023; Cui et al., 2015). As the dsRBM is situated in the CTD, it is retained in N*^M210^ (Fig 1B). To test if N*^M210^ binds to dsRNA, cells expressing N^FL^, N*^M210^, or an empty vector (EV) control were lysed and subjected to a pulldown using streptavidin beads conjugated to biotinylated poly I:C, a synthetic dsRNA mimic. Although both N*^M210^ and N^FL^ co-precipitated with dsRNA, there was 6-fold more N*^M210^ in the eluate compared to N^FL^ (Fig 3A & B). As a control, we mutated the dsRBM of N*^M210^ (K257A/K261 in N^FL^) to generate N*^M210-ΔdsRBM^. No N*^M210-ΔdsRBM^ was detected in the eluate after precipitation.

**Figure 3.**
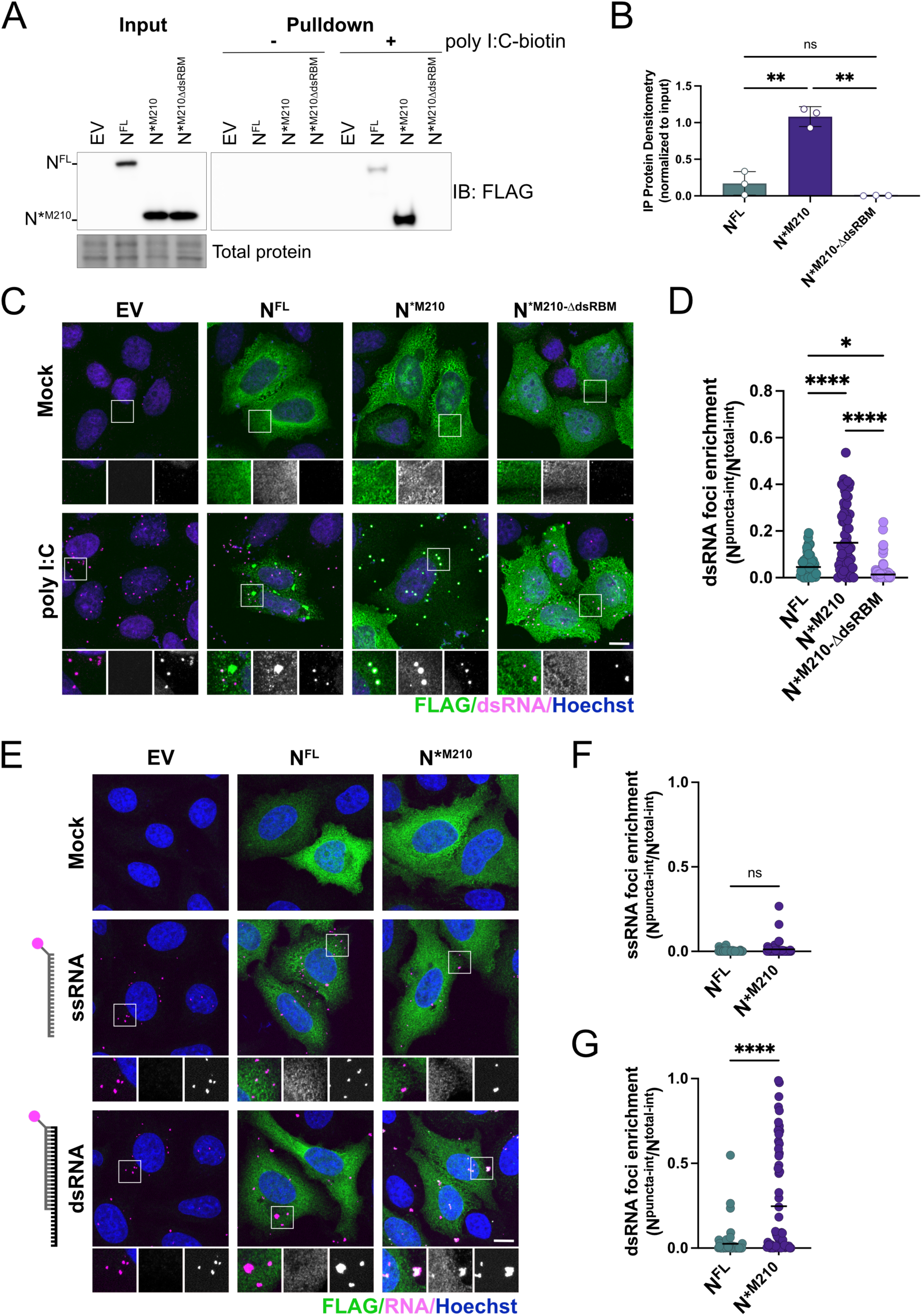
N*^M210^ is a superior dsRNA binding protein compared to N^FL^. **A-B.** EV, N^FL^, N*^M210^, or N*^M210-ΔdsRBM^-transfected HEK293T cells were lysed and subjected to a pulldown using streptavidin beads conjugated to biotinylated poly I:C, or unconjugated beads. Eluted proteins were resolved by SDS-PAGE and immunoblotting. Protein abundance after pulldown was enumerated by densitometry and normalized to each respective input control. These data represent three independent biological replicates (*n=3*). Statistics were performed using a one-way ANOVA with Dunnett’s post-hoc analysis, (**, *p* < 0.0021), mean; standard deviation. **C.** HeLa cells were transfected with FLAG-tagged N^FL^, N*^M210^, or N*^M210-ΔdsRBM^, or an empty vector (EV) control. Internal methionine residues were mutated to ensure only the indicated proteoform of N was expressed ([N^FL^; M210I and M234V], [N*^M210^ and N*^M210-ΔdsRBM^; M234V]). 24 hours post-transfection, cells were transfected with 0.5 µg high molecular weight poly I:C or mock transfected. Three hours after poly I:C treatment, cells were fixed and immunostained with the J2 antibody (dsRNA; Alexa 647) and FLAG antibody (N proteoform; Alexa 488). Nuclei were stained with Hoechst. A maximum intensity projection is presented here. One representative experiment of three independent replicates is shown. Scale bar = 10 µm. **D.** Enrichment of N proteoforms with dsRNA was determined using CellProfiler using confocal images. Enrichment was measured by determining the proportion of N^FL^/N*^M210^ that is cytoplasmic versus co-localized with dsRNA foci. The mean integrated intensity of the N proteoform overlapping with dsRNA foci was divided by the mean integrated intensity of the respective N proteoform in the entire cytoplasm. Each data point represents a single cell. These data represent three independent biological replicates (*n*=3) with >18 cells measured per condition, per replicate. Statistics were performed using a Kruskal-Wallis *H* test with Dunn’s correction (**, *p* < 0.0021, ****, *p* < 0.0001), mean. **E.** HeLa cells were transfected with FLAG-tagged N^FL^, N*^M210^ or EV as in A. 24 hours post-transfection, cells were transfected with 0.9 µg of fluorescein-labelled (18 bp) ssRNA or dsRNA probes (18 bp duplex, 20 bp overhang). Three hours later, cells were fixed and immunostained with the FLAG antibody (N proteoform; Alexa 647). Nuclei were stained with Hoechst. A maximum intensity projection is presented here. One representative experiment of three independent replicates is shown. Scale bar = 10 µm. **F-G.** Enrichment of N proteoforms with RNA probes as in D. These data represent three independent biological replicates (*n*=3) with 20 cells measured per condition, per replicate. Statistics were performed using a Mann-Whitney test (****, *p* <0.0001), mean.

To determine if N*^M210^ sequesters dsRNA in cells, N^FL^, N*^M210^, or EV-expressing HeLa cells were transfected with poly I:C. Immunostaining revealed that poly I:C transfection induced intracellular dsRNA foci (Fig 3C). In mock transfected cells, both N^FL^ and N*^M210^ were diffusely localized throughout the cytoplasm. Three hours after poly I:C treatment, the majority of N^FL^ became concentrated in large perinuclear granules that were spatially distinct from dsRNA foci. A small proportion of N^FL^ did co-localize with dsRNA foci, likely via dsRNA binding. By contrast, N*^M210^ was largely re-localized to dsRNA foci upon poly I:C treatment, suggesting superior dsRNA binding activity compared to N^FL^ (Fig 3C). Quantification of these data illustrates the differential enrichment of N*^M210^ and N^FL^ in dsRNA foci, with an average of 31.5 percent of the total N*^M210^ signal co-localizing with dsRNA, versus 6.4 percent for N^FL^ (Fig 3D). N*^M210-ΔdsRBM^ remained diffuse in the cytoplasm following poly I:C treatment (Fig 3C), indicating that the dsRNA sequestration effect requires this domain. We also confirmed the differential dsRNA sequestration of N^FL^ and N*^M210^ in A549 cells (Fig S3A & B). N*^M210^ also readily sequestered dsRNA upon treatment with low molecular weight (LMW) poly I:C and poly A:U dsRNA, which is more reminiscent of AU-rich CoV RNAs (Fig S4A & B) (Fumagalli et al., 2023). To determine if N*^M210^ has a greater propensity to sequester dsRNA versus single-stranded RNA (ssRNA), N^FL^, N*^M210^, or EV-expressing cells were transfected with a fluorescently labeled ssRNA probe (18 bp) or a dsRNA probe (18 bp duplexed; Fig 3E). N*^M210^ only significantly co-localized with the dsRNA-containing foci, but did not colocalize with ssRNA-containing foci, whereas N^FL^ failed to colocalize with either ss or dsRNA foci (Fig 3F & G).

If both N^FL^ and N*^M210^ contain the dsRBM, we wondered why N*^M210^ was the superior dsRNA binding protein. One explanation is that the amino terminus of N^FL^, which is absent in N*^M210^, negatively regulates the interaction with dsRNA. Phosphorylation of N^FL^ in the SR region has been shown to reduce RNA binding by masking the positive charge of the RNA-binding domains (Botova et al., 2024; C. Wu et al., 2021; J. Wu et al., 2022). The residues necessary for N^FL^ phosphorylation priming, S188 and S206 (C. Wu et al., 2021; Yaron et al., 2022), are upstream from M210, and therefore are not retained in N*^M210^. To determine if N^FL^ phosphorylation can reduce the interaction with dsRNA, thus explaining the enhanced dsRNA-binding capability of N*^M210^ relative to N^FL^, we generated a hypo-phosphorylated N mutant (N^FL-S188A/S206A^) and measured its ability to interact with dsRNA. Contrary to the hypothesis, we found that N^FL-S188A/S206A^ displayed reduced dsRNA foci enrichment compared to N^FL^ (Fig S4C & D). These data show that phosphorylation of N^FL^ in the SR region is not responsible for its inferior ability to sequester dsRNA relative to N*^M210^.

### N*^M210^ can interact with N^FL^, which increases N^FL^ dsRNA sequestration capacity

During infection or over-expression, N^FL^ forms homodimers via a dimerization motif in the CTD and higher order oligomers via a leucine-rich helix in the linker (Peng et al., 2020; Q. Ye et al., 2020; Yu et al., 2005). Since N*^M210^ retains these motifs, we tested if N*^M210^ can interact with N^FL^. We co-expressed N*^M210^-FLAG and N^FL-WT^-HA in HEK293T cells and conducted co-immunoprecipitation assays. We found that N^FL^ readily co-precipitated with N*^M210^ (Fig 4A). The expression construct for N^FL-WT^ contains residues M210 and M234 and produces low levels of N*^M210^ and N*^M234^ in addition to N^FL^ (Fig S1B). Because these N* proteoforms also co-precipitated with N*^M210^, these data reveal that N*^M210^ can interact with N*^M234^ and N*^M210^ proteins, forming homo- and heterodimers (Fig 4B). Co-precipitation of all dimeric combinations (N^FL^-N*^M210^, N*^M210^-N*^M210^, and N*^M210^-N*^M234^) was substantially reduced upon RNase A treatment, suggesting that these interactions are enhanced by the presence of RNA (Fig 4A).

**Figure 4.**
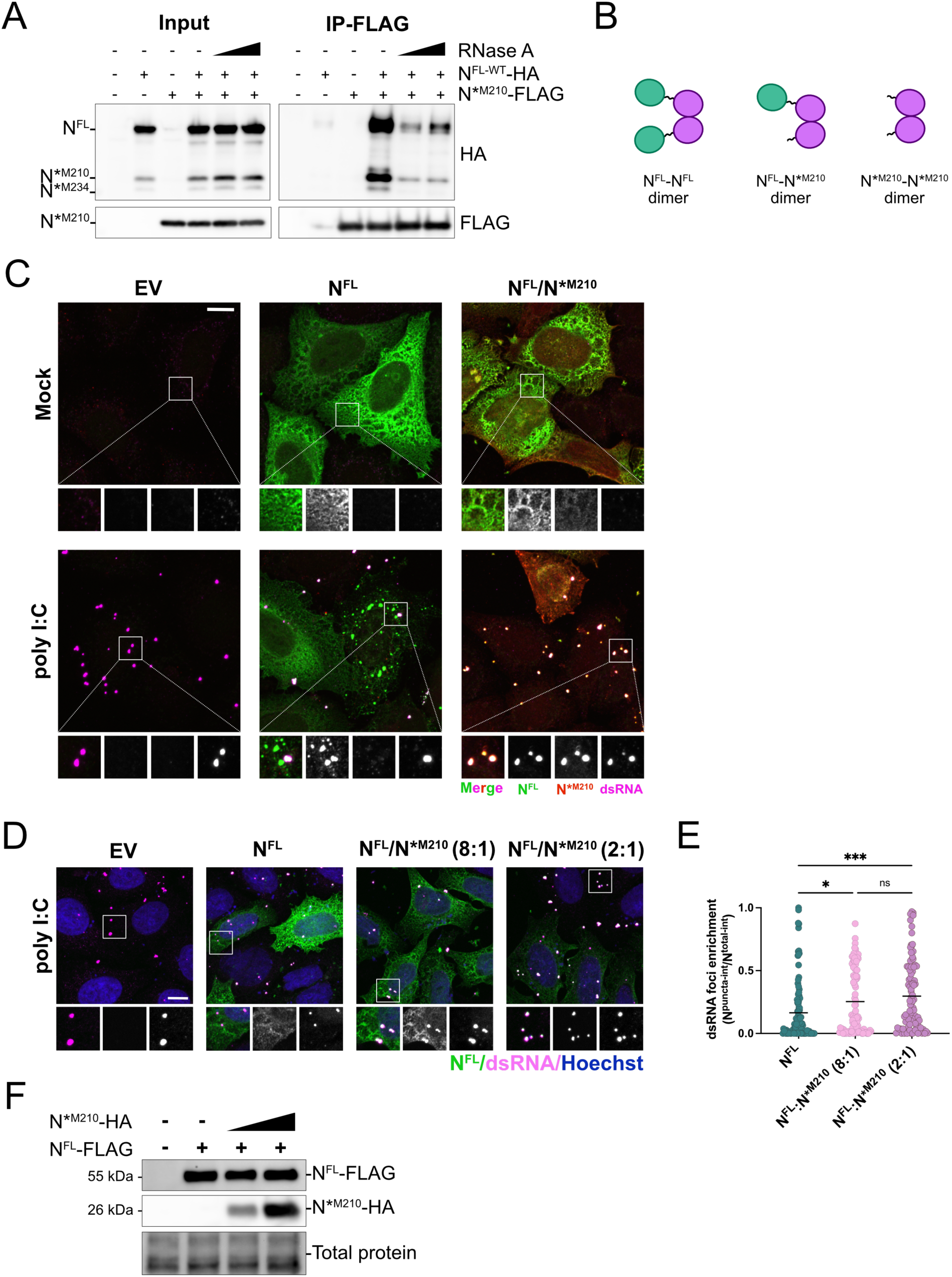
N*^M210^ interacts with N^FL^ to increase N^FL^ co-localization with dsRNA. **A.** HEK293T cells were co-transfected with HA-tagged N^FL-WT^, with intact downstream methionines (M210 and M234), and FLAG-tagged N*^M210^. 24 hours post-transfection cells were lysed, treated with 10 or 50 µg/mL RNase A, and incubated with anti-FLAG antibody overnight at 4°C. Immunoprecipitation was performed using magnetic Dynabeads and samples were resolved by SDS-PAGE and immunoblotted with anti-FLAG and anti-HA antibodies. One of three independent experiments shown. **B.** Schematic representation of N proteoform homo and heterodimers based on A. **C.** HeLa cells expressing N^FL^-FLAG alone or co-expressing N^FL^-FLAG and N*^M210^-HA were transfected with 0.5 µg poly I:C. Three hours post transfection, cells were fixed and immunostained with the FLAG antibody (N^FL^; Alexa 488), the HA antibody (N*^M210^; Alexa 405), and the J2 antibody (dsRNA; Alexa 555). A maximum intensity projection (MIP) is presented here. Scale bar = 10 µm. **D.** HeLa cells expressing N^FL^-FLAG alone or co-expressing N^FL^-FLAG and N*^M210^-HA at an 8:1 or 2:1 N^FL^:N*^M210^ ratio were transfected with 0.5 µg poly I:C. Three hours post transfection, cells were fixed and immunostained with the FLAG antibody (N^FL^; Alexa 488) and the J2 antibody (dsRNA; Alexa 647). Nuclei were stained with Hoechst. A maximum intensity projection (MIP) is presented here. Scale bar = 10 µm. **E.** Enrichment of N^FL^ with dsRNA from D. was calculated as in Fig 3B. These data represent three independent biological replicates (*n*=3) with 37 cells measured per condition, per replicate. Statistics were performed using a Kruskal-Wallis *H* test with Dunn’s correction (****, *p* < 0.0001), mean. **F.** Protein lysate reserved from D was resolved by SDS-PAGE by immunoblotting to validate protein expression.

Given that during an infection, N^FL^ and N*^M210^ are co-expressed in the same cell at the same time (Fig S1C) and that N*^M210^ can interact with N^FL^, we wondered if N*^M210^ can alter the behavior of N^FL^. To test this, we co-expressed N^FL^-HA and N*^M210^-FLAG in HeLa cells, then exposed these cells to poly I:C. We found that upon exposure to dsRNA, N^FL^:N*^M210^ complexes predominantly localized with dsRNA, whereas complexes formed of N^FL^ alone formed distinct, dsRNA-negative foci (Fig 4C). Because N*^M210^ is less abundantly produced than N^FL^ during infection (Fig 1F & G), we tested the relative amount of N*^M210^ required to re-localize N^FL^ to dsRNA foci. We found that co-expression of N*^M210^ increased the propensity for N^FL^ to co-localize with dsRNA foci in a dose-dependent manner. Transfection of an 8:1 ratio of N^FL^:N*^M210^ plasmids was sufficient to induce N^FL^ co-localization with dsRNA; however, N^FL^-dsRNA co-localization was further enhanced after transfection of plasmids at a 2:1 ratio of N^FL^:N*^M210^ (Fig 4D-F). These data show that N*^M210^ and N^FL^ physically interact, an event that alters the behavior and localization of N^FL^.

### N*^M210^ is sufficient to block dsRNA-induced antiviral immune responses

The presence of dsRNA is a telltale sign of a viral infection. As such, dsRNA is a potent inducer of various antiviral immune responses (Y. G. Chen & Hur, 2022; Hur, 2019). There are three primary dsRNA-induced response axes: i) protein kinase R (PKR) binds dsRNA, auto-phosphorylates, then phosphorylates eukaryotic initiation factor 2α, which triggers translational arrest and stress granule formation; ii) RIG-I-like receptors (RLRs) bind dsRNA causing phosphorylation of interferon regulatory factor-3 (IRF3), which induces interferon transcription; and iii) 2’-5’-oligoadenylate synthetase (OAS) binds dsRNA, activating RNase L, which promotes RNA decay and translational arrest (Y. G. Chen & Hur, 2022). N^FL^ has been shown to block all three of these pathways (Aloise et al., 2023; LeBlanc et al., 2023). To test if N*^M210^ is also able to dampen dsRNA-induced immune responses, A549 cells expressing N^FL^, N*^M210^, or EV were treated with poly I:C. Immunoblotting revealed that both N^FL^ and N*^M210^ were sufficient to dampen IRF3 and PKR phosphorylation (Fig 5A-C). Furthermore, N^FL^ and N*^M210^ were sufficient to block *IFNβ* mRNA induction (Fig 5D) and the OAS-RNase L axis, using rRNA decay as a proxy for RNase L activation (Fig 5I & J). The blockade of these three dsRNA-induced responses required the dsRBM of N*^M210^ (Fig 5E-H & J), suggesting N*^M210^ antagonizes these pathways by sequestering dsRNA (Fig 5K).

**Figure 5.**
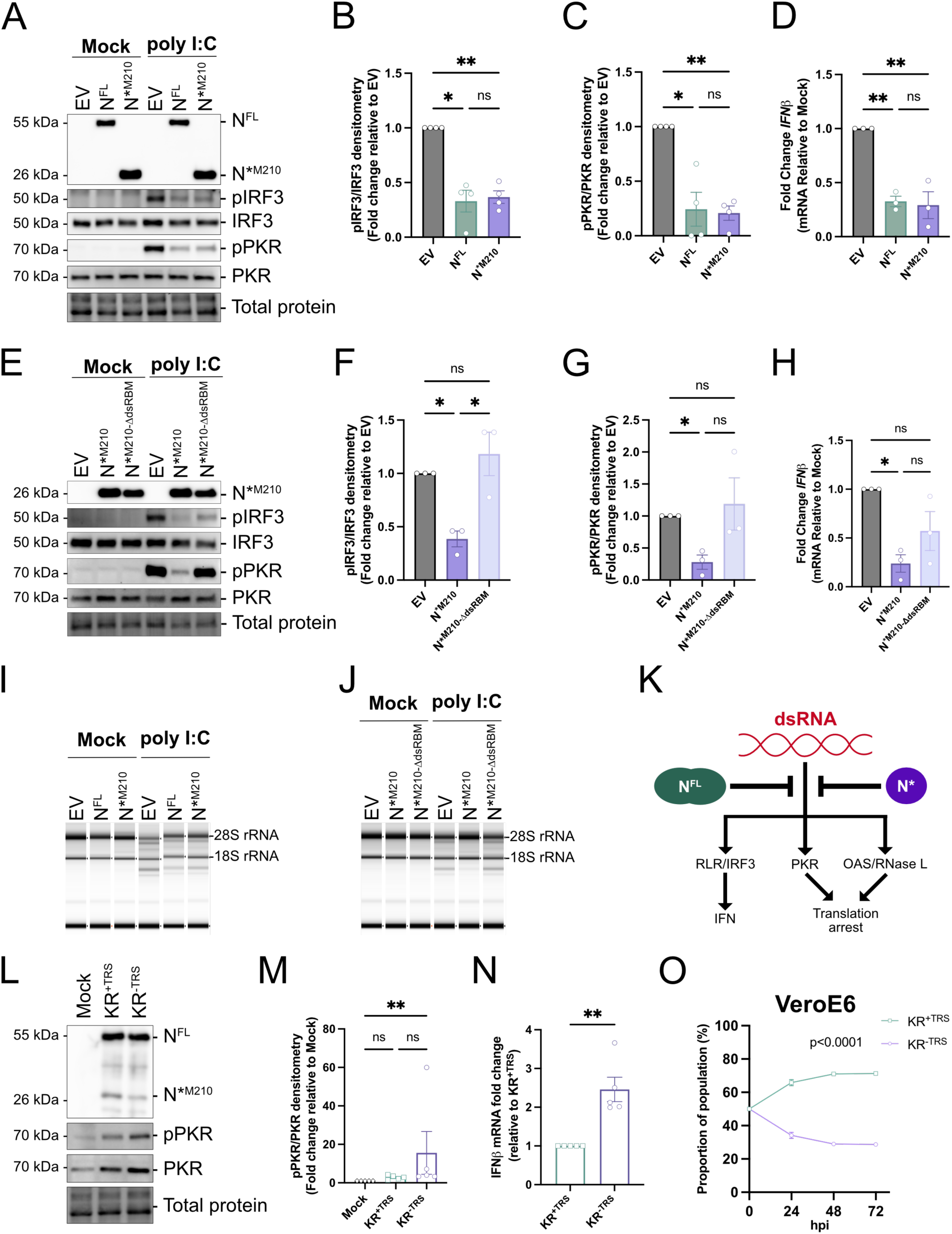
N*^M210^ is sufficient to block cellular dsRNA immune responses. **A.** A549 cells were transduced with recombinant lentiviruses to ectopically express N proteoforms or EV. Internal methionine residues were mutated to ensure only the indicated proteoform of N was expressed ([N^FL^; M210I and M234V], [N*^M210^; M234V]). 96 hours post-transduction, cells were transfected with 1.0 µg high molecular weight poly I:C or mock transfected. Three hours post treatment, protein lysate was harvested and resolved by SDS-PAGE and immunoblotted with antibodies specific for FLAG (N proteoforms), pIRF3, IRF3, pPKR, and PKR. A representative experiment shown. **B-C.** Protein levels from poly I:C-treated samples in A. were quantified by densitometry in ImageLab. pIRF3 and pPKR abundances were measured relative to total IRF and PKR, respectively, and normalized to EV. These data represent four independent biological replicates (*n=4*). Statistics were performed using a one-way ANOVA with Dunnett’s post-hoc analysis. **D.** N proteoforms were ectopically expressed in A549 cells as in A. Cells were treated with 1.0 µg high molecular weight poly I:C or mock transfected. Three hours post poly I:C treatment, intracellular RNA was harvested and *IFN*β RNA abundance was measured by RT-qPCR. These data are represented as fold-change relative to the no poly I:C treatment from three independent biological replicates (*n=3*). Statistics were performed using a one-way ANOVA with Dunnett’s post-hoc analysis. **E.** N proteoforms were ectopically expressed, treated with poly I:C, and subjected to SDS-PAGE and immunoblotting as in A. One of three independent experiments shown. **F-G.** Protein levels from poly I:C-treated samples in E. were quantified by densitometry as in B. and C. These data represent three independent biological replicates (*n*=3). Statistics were performed using a one-way ANOVA with Dunnett’s post-hoc analysis. **H.** N proteoforms were ectopically expressed in A549 cells as in E and treated with poly I:C. Three hours post poly I:C treatment, intracellular RNA was harvested and *IFN*β RNA abundance was measured by RT-qPCR. These data are represented as fold-change relative to the no poly I:C treatment from three independent biological replicates (*n*=3). Statistics were performed using a one-way ANOVA with Dunnett’s post-hoc analysis. **I-J.** RNA lysates from D and H were subjected to automated electrophoresis using an Agilent 4200 TapeStation system to assess rRNA integrity. One representative of three independent experiments shown (*n=3*). **K.** Schematic representation of N^FL^/N*^M210^ inhibition of dsRNA-induced responses. **L.** Primary HUVEC^ACE2^ cells were infected with KR^+TRS^ or KR^-TRS^ recombinant SARS-CoV-2 viruses (MOI of 4), or mock infected. 24 hours post-infection, protein lysate was harvested and resolved by SDS-PAGE and immunoblotted with antibodies specific for N, pPKR, and PKR. A representative experiment shown. **M.** Protein levels from rSARS-CoV-2-infected cells from K. were quantified as in B. PKR levels and relative pPKR abundances were measured and normalized to EV. These data are represented as fold-change relative to mock from five independent biological replicates (*n=5*). Statistics were performed using a Friedman test (non-normal deviation), (*, *p* < 0.05). **N.** HUCEC^ACE2^ cells were infected with KR^+TRS^ or KR^-TRS^, as in K. 24 hours post-infection, intracellular RNA was harvested and *IFN*β RNA abundance was measured by RT-qPCR. Data are represented as fold-change relative to KR^+TRS^. These data represent five independent biological replicates (*n=5*). Statistics were performed using a ratio paired T-test, (**, *p* < 0.0021). **O.** VeroE6 cells were coinfected with equal infectious titers of KR^+TRS^ and KR^-TRS^ recombinant SARS-CoV-2 viruses to achieve a total MOI of 0.01. Time 0 represents the inferred proportions of each recombinant based on infectious titer input. Intracellular RNA was harvested at the indicated times post-infection and subjected to probe-based RT-qPCR to differentiate recombinant virus abundance. These data represent three independent biological replicates (*n=3*). Statistics were performed using a simple linear regression.

To determine if N*^M210^ could impact immune activation in the context of an infection, we infected primary HUVEC^ACE2^ cells with KR^+TRS^ or KR^-TRS^. Both viruses induced an upregulation in total PKR and pPKR levels; however, pPKR levels were slightly higher in cells infected with KR^-TRS^ (Fig 5L & M). Both KR^+TRS^ and KR^-TRS^ increased *IFNβ* abundance, yet KR^-TRS^-infected cells consistently produced more *IFNβ* mRNA than KR^+TRS^-infected cells (Fig 5N). Because N*^M210^ potently blocks IFN induction, we wondered if the KR^+TRS^ virus outcompetes KR^-TRS^ because N*^M210^ antagonizes IFN responses. To test this, we conducted co-infection competition assays in VeroE6 cells, which are unable to produce type I IFNs (Emeny & Morgan, 1979). We reasoned that if the replication advantage of KR^+TRS^ is exclusively due to a superior ability to block IFN, we would expect KR^+TRS^ and KR^-TRS^ to have equal fitness in VeroE6 cells. However, we observed that even in IFN-deficient cells, KR^+TRS^ outcompeted KR^-TRS^, suggesting that the fitness difference between KR^+TRS^ and KR^-TRS^ is not solely from N*^M210^-mediated IFN suppression (Fig 5O). There are two important interpretations from these data: i) N*^M210^ potently antagonizes dsRNA-induced antiviral responses likely via dsRNA sequestration, and ii) N*^M210^ can act to promote viral replication using a mechanism that is independent of IFN responses.

### N*^M210^ inhibits cellular ribonucleoprotein granules via the dsRBM

Stress granules (SGs) are thought to have antiviral activity, providing an obstacle for a productive viral infection (Dolliver et al., 2022; McCormick & Khaperskyy, 2017; Ruggieri et al., 2012; W. Yang et al., 2019). SGs are cytoplasmic granules that form after translation initiation blockage. This can occur by phosphorylation of eIF2α by stress-activated kinases, including PKR from dsRNA/viral infection and heme-regulated inhibitor kinase (HRI) from oxidative stress (McCormick & Khaperskyy, 2017; Protter & Parker, 2016). Translationally repressed mRNAs coalesce with RNA-binding proteins, such as Ras GTPase-activating protein-binding protein 1 (G3BP1) and T cell intracellular antigen-1 (TIAR), resulting in SG formation. Recently, the Parker and Burke labs have shown that dsRNA induces a G3BP1-positive SG-like condensate called an RNase-L dependent body (RLB) (Burke et al., 2019, 2020). We acknowledge SG diversity, however for simplicity, here we referred to all G3BP1-positive foci as SGs. The SARS-CoV-2 N^FL^ protein has been reported to co-localize with SGs and block their formation (Aloise et al., 2023; Biswal et al., 2022; Luo et al., 2021; Zheng et al., 2021). To test if N*^M210^ can also block SGs, HeLa cells expressing N*^M210^, N^FL^, or EV were treated with poly I:C, which induces SG formation via PKR. Using immunofluorescence microscopy with G3BP1 as a marker for SGs, we found that both N^FL^ and N^FL-ΔdsRBM^ readily localized to SGs, consistent with prior reports (Biswal et al., 2022; Z. Yang et al., 2024). Both N*^M210^ and N*^M210-ΔdsRBM^ did not co-stain with SGs and thus, must lack this co-localization motif (Fig 6A). Instead, N*^M210^ formed G3BP1-negative puncta, reminiscent of dsRNA foci from Fig 3C, whereas N*^M210-ΔdsRBM^ remained diffuse following poly I:C treatment. We found that N^FL^ reduced SG formation, but this effect was marginal, especially relative to N*^M210^ which potently blocked SGs using a mechanism dependent on the dsRBM (Fig 6A & B).

**Figure 6.**
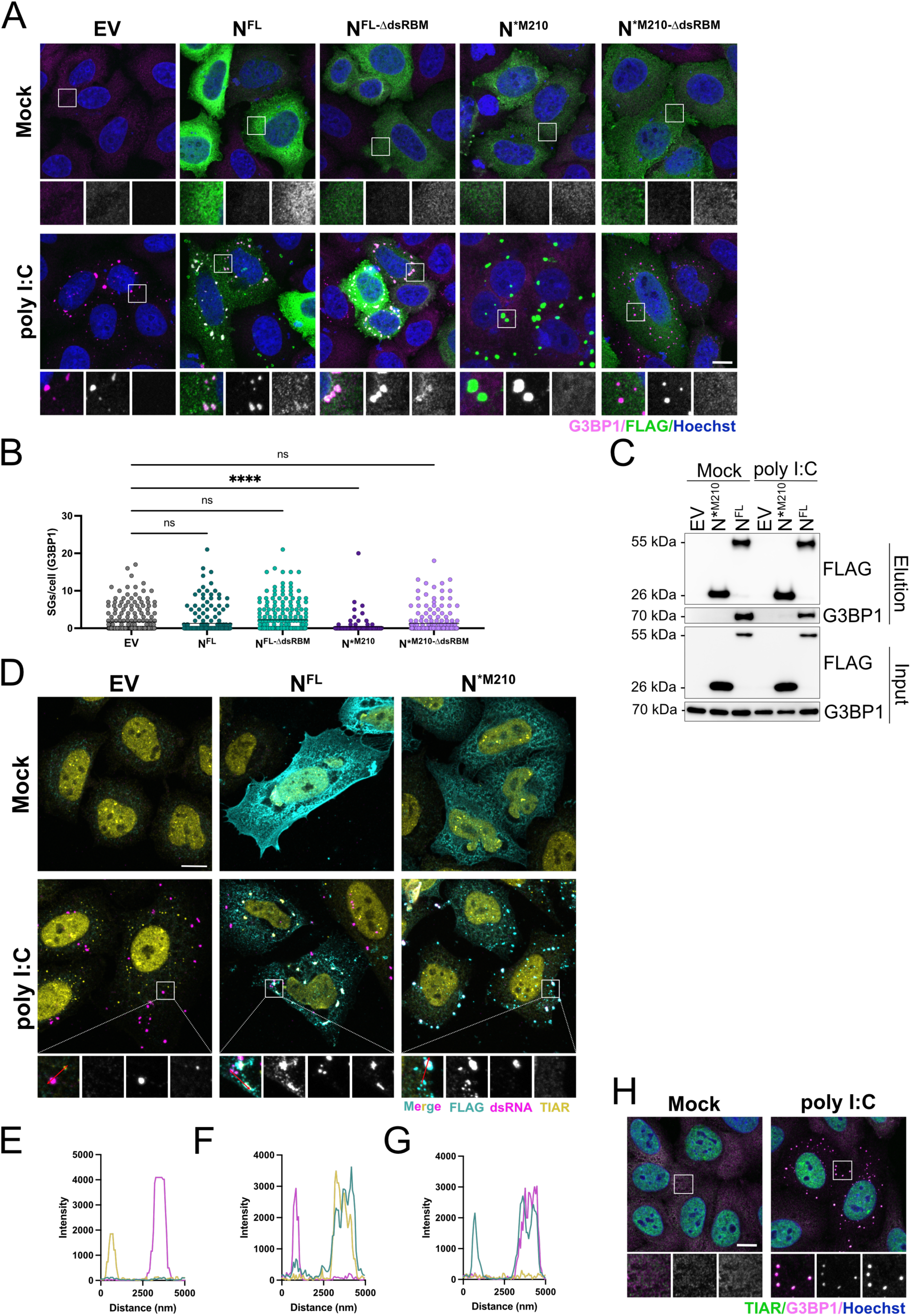
N*^M210^ blocks dsRNA-induced SGs by dsRNA binding. **A.** HeLa cells were transfected with FLAG-tagged N^FL^, N^FL-ΔdsRBM^, N*^M210^, or N*^M210-ΔdsRBM^, or an empty vector (EV) control. Internal methionine residues were mutated to ensure only the indicated proteoform of N was expressed ([N^FL^ and N^FL-ΔdsRBM^; M210I and M234V], [N*^M210^ and N*^M210-ΔdsRBM^; M234V]). 24 hours post-transfection, cells were transfected with 0.5 µg poly I:C or mock transfected. Three hours after poly I:C treatment, cells were fixed and immunostained with a G3BP1 antibody (SGs; Alexa 647) and FLAG antibody (N proteoform; Alexa 488). Nuclei were stained with Hoechst. A maximum intensity projection is presented here. Scale bar = 10 µm. **B.** SGs were quantified using CellProfiler by measuring G3BP1 puncta in N-expressing cells (thresholded by FLAG staining) or EV-transfected cells. These data represent three independent biological replicates (*n* = 3) with >50 cells measured per condition, per replicate. Each datapoint represents a single cell. Statistics were performed using a Kruskal-Wallis *H* test with Dunn’s correction (****, *p* < 0.0001, mean). **C.** HEK293T cells were transfected with pcDNA-N^FL^-FLAG (M210I, M234V), pcDNA-N*^M210^-FLAG (M234V), or an empty vector (EV) control. 48 hours post-transfection, cells were transfected with 10 µg of high molecular weight poly I:C. Three hours post poly I:C treatment, cells were lysed and incubated with anti-FLAG antibody overnight at 4°C. Immunoprecipitation was performed using magnetic Dynabeads (Thermo-Fisher) and sample were resolved by SDS-PAGE and immunoblotted with anti-FLAG antibody and anti-G3BP1 antibody. One of three independent experiments shown. **D.** HeLa cells were transfected with N^FL^, N*^M210^, or EV, and treated with 0.5 µg poly I:C as in A. Three hours after poly I:C treatment, cells were fixed and immunostained with antibodies specific for TIAR (SGs; Alexa405), J2 (dsRNA; Alexa647), and FLAG (N proteoform; Alexa488). A maximum intensity projection (MIP) is presented here. Scale bar = 10 µm. **E-G.** Intensity histograms were obtained in Zen Blue. Line was drawn through confocal images from E (inset, red line) and intensity maps are depicted. **H.** HeLa cells were transfected with 0.5 µg poly I:C. Three hours after poly I:C treatment, cells were fixed and immunostained with antibodies specific for TIAR (Alexa 488) and G3BP1 (Alexa 647). Nuclei were stained with Hoechst. A maximum intensity projection (MIP) is presented here. Scale bar = 10 µm.

Previous work has shown that N^FL^ inhibits SG formation by sequestering G3BP1 via the φxF motif in the amino-terminus of N^FL^ (Fig 1B; Biswal et al., 2022; Z. Yang et al., 2024). We found that unlike N^FL^, N*^M210^ did not co-immunoprecipitate with G3BP1 and co-localization with G3BP1 was not impacted by poly I:C transfection (Fig 6C). By co-staining for dsRNA, FLAG, and TIAR, an alternative SG marker, we confirmed that poly I:C treatment induces N^FL^ localization to SGs and N*^M210^ localization to dsRNA foci (Fig 6D-G) and that TIAR does co-localize to G3BP1-positive SGs following poly I:C transfection (Fig 6H). Collectively, these data suggest that N*^M210^ strongly blocks dsRNA-induced SGs, using a mechanism that is independent of G3BP1 binding or co-localization, but is dependent on the dsRBM.

Given that N*^M210^ blocks poly I:C-induced SGs, and that requires the dsRBM, we hypothesized that N*^M210^ inhibits SGs by sequestering dsRNA upstream of PKR activation. We reasoned that N*^M210^ would not block SGs induced by alternative stressors, such as sodium arsenite (SA) that induces SG formation by phosphorylation of HRI (Lu et al., 2001). To test this, we treated cells expressing N*^M210+/-dsRBM^ or N^FL+/-dsRBM^ with SA. We observed that both N^FL^ and N^FL-ΔdsRBM^ co-localized with SA-induced SGs whereas N*^M210^ and N*^M210-ΔdsRBM^ remained diffuse in the cytoplasm (Fig 7A & B). N^FL^ blocked SG formation following SA treatment; however, this was largely dependent on the dsRBM, as N^FL-ΔdsRBM^ did not significantly reduced SGs. We also found that formation of SA-induced SGs was inhibited by N*^M210^, but not N*^M210-ΔdsRBM^. There are two important interpretations from these data: i) N*^M210^ does not only block PKR-induced SGs by sequestering dsRNA, but instead limits SG formation from various inducers and ii) the dsRNA-binding activity of N*^M210^ is required to antagonize SG formation, even when SG formation is induced via a dsRNA-independent pathway.

**Figure 7.**
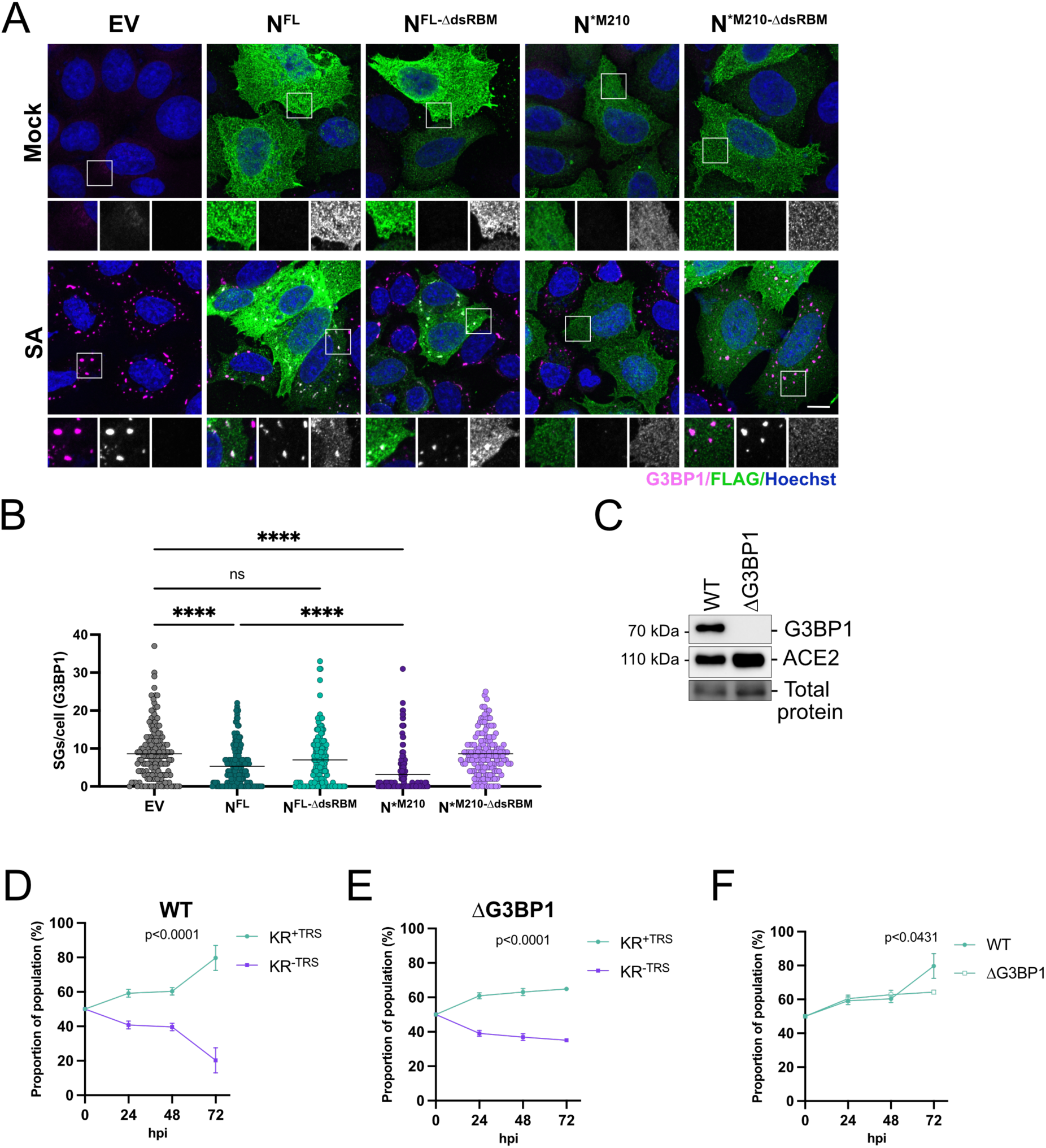
The virus fitness advantage provided by N*^M210^ is tempered in ΔG3BP1 cells. **A.** HeLa cells were transfected with FLAG-tagged N^FL^, N^FL-ΔdsRBM^, N*^M210^, or N*^M210-ΔdsRBM^, or an empty vector (EV) control. Internal methionine residues were mutated to ensure only the indicated proteoform of N was expressed ([N^FL^ and N^FL-ΔdsRBM^; M210I and M234V], [N*^M210^ and N*^M210-ΔdsRBM^; M234V]). 24 hours post-transfection, cells were treated with 500 µM sodium arsenite for 1 hour before the cells were fixed and immunostained with a G3BP1 antibody (SGs; Alexa 647) and FLAG antibody (N proteoform; Alexa 488). Nuclei were stained with Hoechst. A maximum intensity projection is presented here. Scale bar = 10 µm. **B.** SGs were quantified as in Fig 6B. These data represent four independent biological replicates (*n*=4) with 40 cells measured per condition, per replicate. Each datapoint represents a single cell. Statistics were performed using a Kruskal-Wallis *H* test with Dunn’s correction (****, *p* < 0.0001, mean). **C.** Protein lysates from wild-type and G3BP1 knockout HEK293A^ACE2^ (ΔG3BP1) cells were harvested and resolved by SDS-PAGE and immunoblotted with anti-G3BP1 and anti-ACE2 antibodies. **D-E.** Wildtype (WT) or G3BP1 knockout (ΔG3BP1) HEK293^ACE2^ cells were coinfected with equal infectious titers of KR^+TRS^ and KR^-TRS^ recombinant SARS-CoV-2 viruses to achieve a total MOI of 0.01. Time 0 represents the inferred proportions of each recombinant based on infectious titer input. Intracellular RNA was harvested at the indicated times post-infection and subjected to probe-based RT-qPCR to differentiate recombinant virus abundance. These data represent four independent biological replicates (*n*=4). Statistics were performed using a simple linear regression. **G.** The relative proportions of the KR^+TRS^ virus in WT and ΔG3BP1 cells(from the coinfection assays from D. and E.) were overlayed to compare differences in fitness. Differences in slopes were compared using a simple linear regression.

In addition to blocking SGs, N^FL^ also induces processing body (P-body) disassembly (Kleer et al., 2022). P-bodies are also membraneless granules, but unlike SGs, P-bodies are constitutively present in the cytoplasm where they act as sites of RNA repression and decay (Blake et al., 2024; Hubstenberger et al., 2017; Sheth & Parker, 2003; Teixeira et al., 2005). P-bodies are hypothesized to have antiviral activity, therefore many viruses, including SARS-CoV-2, trigger P-body disassembly during infection (Corcoran et al., 2012; Gaete-Argel et al., 2019; Kleer et al., 2022). Unlike SGs, P-bodies are not induced by dsRNA (Burke et al., 2020). To test if N*^M210^ is sufficient to disassemble P-bodies, HUVEC cells expressing N^FL^ or N*^M210^ with or without the dsRBM were fixed and immunostained (Fig S5A). P-bodies were quantified by measuring Hedls, a resident P-body protein. Both N^FL^ and N*^M210^ decreased P-body numbers and both proteoforms require an intact dsRBM to mediate P-body disassembly (Fig S5B). Collectively, these data suggest that N*^M210^ potently blocks SGs and P-bodies using a mechanism that relies on its dsRBM.

If SGs are a hurdle for a successful virus infection and N*^M210^ blocks SG formation, we wondered if the difference in virus fitness between KR^+TRS^ and KR^-TRS^ could be explained by the ability of N*^M210^ to block SG formation. To directly test if KR^+TRS^ outcompetes KR^-TRS^ because of an enhanced ability to inhibit SGs, we conducted co-infection competition assays in either wild-type (WT) or G3BP1 knockout (ΔG3BP1) HEK293A^ACE2^ cells, the latter of which have impaired SG formation (Tsai et al., 2016) (Fig 7C). Consistent with A549^ACE2^ and HUVEC^ACE2^ cells, KR^+TRS^ readily outcompeted KR^-TRS^ in WT HEK293A^ACE2^ cells, as by 72 hpi KR^+TRS^ comprised 80 percent of the virus population (Fig 7D). In ΔG3BP1 HEK293A^ACE2^ cells, KR^+TRS^ still outcompeted KR^-TRS^, however the fitness advantage of KR^+TRS^ was significantly diminished in knockout cells compared to WT cells, with KR^+TRS^ plateauing at 60 percent dominance (Fig 7D-F). These data suggest that in cells unable to form antiviral SGs, these viruses have converged in terms of fitness, signifying that increased N*^M210^ production by KR^+TRS^ confers a virus fitness advantage, in part, by more efficiently antagonizing SGs.

Taken together, we show that dominant SARS-CoV-2 variants have evolved to produce the truncated N proteoform (N*^M210^) in greater abundance, where increased N*^M210^ production increases SARS-CoV-2 fitness by antagonizing of multiple arms of the antiviral response. However, competition assays in SG- and IFN-incompetent cells suggests that N*^M210^-induced SG blockade plays the dominant role in enabling viruses with increased N*^M210^ to outcompete wild-type viruses.

## DISCUSSION

The host-pathogen relationship during a viral infection can be likened to a molecular game of chess, where each offensive move is rapidly countered until one side ultimately wins. However rather than pushing pieces on a board, the ‘moves’ are genetic acquisitions, which gives either the virus or the cell the edge to succeed. There are two ways mutations can improve virus fitness: increase the efficiency of the virus replication cycle; or decrease the effectiveness of the host antiviral response. Here, we show that a genetic acquisition by SARS-CoV-2 enhances the synthesis of a truncated nucleocapsid proteoform (N*^M210^) to give select variants a competitive edge by antagonizing multiple branches of the cellular antiviral response. Our three major findings are as follows: i) SARS-CoV-2 has acquired mutations to increase the production of N*^M210^; ii) N*^M210^ is a double-stranded RNA binding protein that diminishes the interferon response and blocks antiviral granules; and iii) the production of N*^M210^ provides a virus fitness advantage, in part by conferring resistance to antiviral stress granules.

### SARS-CoV-2 produces truncated N proteoforms

Viruses have genomes that are magnitudes smaller than mammalian genomes, yet despite this, viruses still efficiently hijack and undermine cellular defences. One of the ways viruses circumvent this imbalance is by maximizing their genetic economy. For example, many viruses produce truncated versions of their proteins, which can alter protein function without requiring additional genetic space (Baggio et al., 2021; Bostedt et al., 2024; Corcoran et al., 2009; Reuven et al., 2004). Here, we show that SARS-CoV-2 produces multiple truncated versions of the N protein, including N*^M210^, N*^M234^, and N*^37kDa^. Our results show that internal translation initiation from methionine codons in positions 210 and 234 produces N* proteoforms N*^M210^, and N*^M234^ (Fig 1I & S1A & B). Although the N gene also contains a putative in-frame start codon at M101, we did not detect this N* proteoform after over-expression (Fig 1I). This could suggest that this AUG is suboptimal for translational initiation, or that the protein product is unstable. We suspect that N*^37kDa^ is a cleavage product that either requires other viral proteins or the induction of an antiviral state to form as it is only detectable after immunoblotting of infected cell lysate and not after ectopic expression of N^FL^.

### N*^M210^ is a viral dsRNA binding protein

The N protein contains two structured domains that can bind RNA, the NTD and CTD; the NTD has an affinity for TRS-like ssRNA and the CTD, containing the dsRBM, prefers dsRNAs (Iserman et al., 2020; Roden et al., 2022). Because N*^M210^ contains the entire CTD we expected N*^M210^ to retain dsRNA binding activity. Indeed, we found that N*^M210^ interacts with multiple types of dsRNA, including dsRNA as small as 18 duplexed nucleotides (Fig 3E). We show that both N*^M210^ and N^FL^ more readily localizes to dsRNA compared to ssRNA. We did not observe extensive co-localization between N^FL^ and ssRNA; however, it is possible that the 18 bp ssRNA probe is below the optimal length or is not the ideal sequence (Cubuk et al., 2024; Iserman et al., 2020). Using orthogonal approaches, we show that N*^M210^ has an increased ability to bind dsRNA compared to N^FL^ despite sharing an identical dsRBM (Fig 3, S3, S4).

If both N^FL^ and N*^M210^ share the same dsRBM, why would N*^M210^ have increased ability to interact with dsRNA? One explanation is that the amino-terminus of N^FL^ somehow interferes with dsRNA binding. We found that phosphorylation of the SR region of N^FL^ does not explain its reduced dsRNA binding capacity (Fig S4C-D). Another possible explanation for N*^M210^ superior dsRNA binding ability is phase separation. N^FL^ has been shown to phase separate with RNA impacting RNA-binding (Cubuk et al., 2021; Iserman et al., 2020; Perdikari et al., 2020; Roden et al., 2022); therefore, it is possible that removal of the amino-terminus alters the phase separation properties of N*^M210^, allowing for a more stable interaction with dsRNA.

### N*^M210^ blocks dsRNA-mediated immune responses and antagonizes RNP granules

The N^FL^ protein is a multifunctional protein with various roles during viral replication, including promoting viral RNA synthesis and packaging as well as antagonizing cellular antiviral responses (Aloise et al., 2023; LeBlanc et al., 2023). We found that N*^M210^ represses antiviral pathways by sequestering dsRNA. Although N*^M210^ has an increased ability to interact with dsRNA compared to N^FL^, N*^M210^ was not significantly better at blocking dsRNA-induced responses (Fig 5). This inconsistency is likely a result of an excess of N^FL^/N*^M210^ during over-expression compared to dsRNA, which could mask the potential differences between these two proteoforms.

N^FL^ also inhibits SG formation (Dolliver et al., 2022; Zheng et al., 2021). SGs have been shown to form in response to SARS-CoV-2 infection and several studies have shown that SGs restrict CoV replication (Long et al., 2024). Furthermore, SG inhibition via G3BP1 knockdown/knockout increased SARS-CoV-2 replication *in vitro* and *in vivo* (Liu et al., 2022; Zheng et al., 2021). Similarly, Dolliver et al (2022) found that over-expression of G3BP1 increased SG formation and correspondingly decreased the replication of the common-cold coronavirus HCoV-OC43. Precisely how SGs exercise their antiviral activity remains unclear; however, Burke et al (2024) found that G3BP1 can antagonize SARS-CoV-2 replication by trapping and repressing SARS-CoV-2 RNAs. In a complementary study, Yang et al (2024) illustrated that without the G3BP1-interacting motif in N, SARS-CoV-2 replication was diminished, supporting a model where the N-G3BP1 interaction is necessary for optimal replication by preventing viral RNA translational repression.

Although factors that influence SG formation and stability remain incompletely understood, G3BP1/2 and RNA are necessary for SG scaffolding (Decker et al., 2022; Guillén-Boixet et al., 2020; Parker et al., 2025; Tourrière et al., 2023). Previous studies have shown that N^FL^ inhibits SGs by two primary mechanisms: i) N^FL^ interacts with G3BP1 via a φxF motif in the N-IDR of N^FL^ (Fig 1B), thus disrupting SG stability/assembly (Huang et al., 2021; Kruse et al., 2021; Luo et al., 2021; Z. Yang et al., 2024); and ii) N^FL^ sequesters dsRNA to block PKR activation and prevents dsRNA-induced SG formation (Aloise et al., 2023; Zheng et al., 2021). We show that N*^M210^ inhibits SGs independent of any G3BP1 interaction but was dependent on the dsRBM. We found that N*^M210^ blocked SGs induced by poly I:C (dsRNA) and sodium arsenite (oxidative stress), the latter of which does not rely on dsRNA to activate PKR (Lu et al., 2001). This finding suggests that N*^M210^ inhibits SGs independent G3BP1 binding and possibly independent of PKR activation. Burke et al (2020) found that dsRNA-induced SGs/RLBs did not require PKR activation to form. This is consistent with a model where N*^M210^ inhibits SGs independent of PKR (Burke et al., 2020). Given that SGs rely on extensive RNA-RNA interactions (Tauber et al., 2020), it is possible that N*^M210^ sequesters cellular RNAs, preventing their condensation, and thus preventing SG formation. This phenomenon has been previously observed by the Parker lab, who showed that eIF4A antagonizes SGs by de-condensing RNAs (Burke et al., 2024). In support of N*^M210^ as an “RNA de-condenser”, we show that N*^M210^ also inhibits P-bodies in a dsRBM-dependent manner, suggesting that N*^M210^ could broadly antagonize RNP granules by interfering with RNA-RNA scaffolding.

In this study, we focused on the functions of N*^M210^ as an immune antagonist; however, N*^M210^ may also participate in other aspects of the viral infectious cycle. Recently, the Morgan and Doudna labs found that N*^M210^ participates in viral packaging (Adly et al., 2023; Syed et al., 2024). In support of this finding, we show that N*^M210^ and other N* proteoforms are found in extracellular viral particles (Fig 1H). To our knowledge, we are the first group to show truncated N proteoforms packaged into authentic SARS-CoV-2 virions, raising the interesting possibility that packaged N*^M210^ could play a role during the early phase of viral infection. Furthermore, Bouhaddou et al (2023) found that N*^M210^ interacts with the RNA polymerase II-associated factor (PAF) complex, suggesting that N*^M210^ could have nuclear localization. In agreement with this, we observe some nuclear localization after ectopic expression of N*^M210^ (Fig 3C), although the biological consequences of N*^M210^-PAF interaction are unknown.

### N*^M210^ production increases virus replication fitness

All SARS-CoV-2 variants we tested produced multiple truncated N proteoforms, however N*^M210^ and N*^M234^ production was upregulated in Alpha, Gamma, and Omicron due to the internal TRS acquisition. This suggests that N* production may be undergoing positive selection due to its importance for SARS-CoV-2 replication in humans. To directly determine if N*^M210^ production impacts virus fitness, we conducted co-infection competition assays and found that the internal TRS, which promotes N*^M210^ synthesis, confers a replication advantage in multiple cell lines and primary human cells (Fig 2, 3, 7 & S2). Previous studies illustrated that the R203K/G204R amino acid substitution conferred a fitness advantage (Johnson et al., 2022; H. Wu et al., 2021). They found that R203K/G204R enhances N^FL^ phosphorylation, phase separation, and antiviral suppression (Johnson et al., 2022; H. Wu et al., 2021; Zhao et al., 2021). However, the R203K/G204R amino acid change also introduces the internal TRS, which upregulates N*^M210^ synthesis, and these studies did not consider the impact of increased N*^M210^ production on viral fitness. We directly answer this question using recombinant viruses to uncouple the role of the new TRS and the role of altered N^FL^ activity due to amino acid changes. We reveal that the TRS, and corresponding upregulation of N*^M210^ production, increases virus fitness independent of amino acid changes, as rSARS-2 KR^+TRS^ outcompeted both the wild-type (WT) and the amino acid-matched KR^-TRS^ viruses. However, the fitness advantage of KR^+TRS^ compared to WT was greater than KR^+TRS^ compared to KR^-TRS^ (Fig 2E, F, H, I). This supports a model whereby the R203K/G204R substitution enhances SARS-CoV-2 fitness in two ways: i) at the amino acid level, by improving N^FL^ antiviral activity and ii) at the nucleotide level, by promoting N*^M210^ production, a protein which potently represses antiviral pathways. We did not observe a fitness difference between the WT and the M210I viruses, suggesting that basal production of N*^M210^ was insufficient to improve virus replication fitness *in vitro*. It is possible that in the context of a viral infection in an animal model, low levels of N*^M210^ produced by internal translation initiation may influence viral fitness.

We show that N*^M210^ antagonism of antiviral responses contributes to the observed fitness advantage, as rSARS2 KR^+TRS^ was superior at blocking IFNβ induction (Fig 5M). Furthermore, we show that N*^M210^ provides a fitness advantage, in part, by antagonizing SGs, as the replication difference between KR^+TRS^ and KR^-TRS^ was significantly diminished in ΔG3BP1 cells compared to WT cells (Fig 7D-F). This finding indicates that N*^M210^ promotes virus fitness in multiple ways, one of which is by antagonizing SGs. During the writing of this manuscript, Mears et al (2025) published a complementary study supporting our finding that the acquisition of the novel TRS in N improves virus fitness and showing that N*^M210^, which they call N.iORF3, impacts fitness, in part, by inhibiting RIG-I activation. However, in this study the mechanism of RIG-I inhibition was not determined (Mears et al., 2025). Our data support and extend their observations by providing a mechanistic explanation for how N*^M210^ antagonizes RIG-I activation, namely by sequestering dsRNA. This work is a significant contribution to the field as it mechanistically illustrates how N*^M210^ antagonizes multiple antiviral pathways, including RIG-I/IFN, P-bodies, and SGs to promote viral replication in human cells.

Before spilling over into the human population, SARS-CoV-2 or its recent ancestors were primarily circulating in horseshoe bats (Andersen et al., 2020), begging the question, why did this TRS mutation arise during human infection and not in bats? We speculate that upon entering the human host, SARS-CoV-2 faced new immune pressures, incentivizing the upregulation of N*^M210^. We are beginning to understand the differences between human and bat interferon responses; however, to our knowledge, the role of RNP granules in antiviral defense has never been investigated in bats. Because bats are reservoirs for many viruses with pandemic potential, it is critical to understand the immunological pressures in bats and humans to better predict and mitigate the emergence of high-risk pathogens.

Here, we characterize a mutation that arose in the SARS-CoV-2 nucleocapsid gene. We show that this mutation enables the synthesis of a new viral sgRNA, increasing the production of a truncated N protein, N*^M210^, which is a potent dsRNA-binding protein capable of antagonizing the cellular antiviral response. Using recombinant SARS-CoV-2 viruses, we show that the upregulation of N*^M210^ increases viral antagonism of the host response, giving these viruses the edge to outcompete other variants.

### Limitations of this study

Because the anti-N antibody epitope is in the CTD, we were only able to detect amino terminally truncated proteoforms. Therefore, all potential carboxy terminally truncated products would not be detected. Furthermore, the co-infection competition assays measure the proportion of a viral species over time. The dominance of a certain species may reflect increased antiviral tolerance/evasion or increased replication kinetics or efficiency. In short, if a viral species can replicate faster, it will likely outcompete its slower counterpart in these co-infection assays. We also consider that the limitation of a co-infection competition assay is that there is potential for cooperative use of N*^M210^ by both recombinants if they co-infect the same cell. In such cases, N*^M210^ produced by KR^+TRS^ could be used to enhance replication of both viral populations; for this reason, these assays are likely to underrepresent the fitness advantage of KR^+TRS^, further highlighting the significance of N*^M210^ in virus replication. Additionally, N*^M210^ evolved in the context of an *in vivo* infection, not cell culture. By using a cell culture model, we are limited to characterizing the intracellular role of N*^M210^. It is possible that *in vivo* N*^M210^ influences both intrinsic and adaptive immune responses.

## MATERIALS AND METHODS

### Cell culture

All cells were grown in a humidified chamber at 37°C, 5% CO_2_, and 20% O_2_. A549 (ATCC), HEK293T (ATCC), HEK293A (ATCC), HEK293A-ΔG3BP1 (a generous gift from Dr. Denys Khaperskyy, Dalhousie University), HeLa (ATCC), VeroE6 (ATCC), and VeroE6^TMPRSS2^ cells (a generous gift from Dr. Michael Joyce and Holly Bandi, University of Alberta) were cultured in DMEM (Thermo-Fisher) supplemented with 10% heat-inactivated fetal bovine serum (Thermo-Fisher) and penicillin (100U/mL), streptomycin (100U/mL), L-glutamine (2mM) (PSQ; Thermo-Fisher). Calu3 (ATCC) cells were cultured in EMEM (ATCC) supplemented with 10% heat-inactivated fetal bovine serum (Thermo-Fisher) and penicillin (100U/mL), streptomycin (100U/mL), L-glutamine (2mM). HUVECs (Lonza) were cultured in EGM2 (Lonza). HUVECs were seeded onto glass coverslips or plastic dishes coated in 0.1% (w/v) porcine gelatin (Sigma) in 1X PBS (Thermo-Fisher). HUVECs were used between passages 5-7.

A549^ACE2^, HEK293A^ACE2^, and HEK293A-ΔG3BP1^ACE2^ cells were generated by lentivirus transduction of ACE2. Cells were cultured for 48-72 hours in 5 µg/mL blasticidin (Thermo-Fisher) to select for ACE2-expressing cells. Following selection, cells were cultured in DMEM with 10% FBS and 1X PSQ. HUVEC^ACE2^ were generated by lentivirus transduction of ACE2 at passage 5, selected with 5 µg/mL blasticidin for 48 h, then used for virus infections at passage 6.

### Plasmids and cloning

SARS-CoV-2 codon-optimized, FLAG-tagged nucleocapsid (pLJM1-N^FL^-FLAG; Kleer et al. 2022) was subcloned into pcDNA3.1 and pLVX-IRES-puro backbones. N truncations (N*) were generated by PCR amplification and restriction digestions using BamHI and EcoRI (NEB) restriction sites. N amino acid substitutions were generated by site-directed mutagenesis of the pcDNA3.1-N^FL^-FLAG or pcDNA3.1-N*^M210^-FLAG plasmids. Methionine substitutions were selected based on conserved residues in related beta-CoV HCoV-OC43. pcDNA3.1-N^FL^-HA and pcDNA3.1-N*^M210^-HA were cloned from pcDNA3.1-N^FL^-FLAG and pcDNA3.1-N*^M210^-FLAG, respectively. To generate standards for qPCR, a 153 bp fragment of ORF1ab was cloned from SARS-CoV-2-infected HUVEC^ACE2^ cells and subcloned into pcDNA3.1 using BamHI and EcoRI. Standards for co-infection competition assays were generated by subcloning fragments from the BACmid (see Methods “recombinant virus cloning and mutagenesis) containing the N open reading frame into pcDNA3.1 using NheI and NotI. All PCRs were performed with Phusion High-Fidelity PCR Master Mix with GC Buffer (New England Biolabs).

### Recombinant virus cloning and mutagenesis

For viral reverse genetics and recombinant virus rescue, we used the infectious cDNA clone bacterial artificial chromosome (BAC) system (a kind gift from Dr. Luis Martinez-Sobrido, Texas Biomedical Research Institute) (Chiem et al., 2021; C. Ye et al., 2020). For N gene mutagenesis, a site-directed mutagenesis overlap extension PCR and Gibson assembly approach was used. For Gibson assembly, the BAC was divided into 4 fragments of 5-6 kb and one larger fragment of 15.4 kb containing the vector backbone. All 5-6 kb fragments were amplified by conventional PCR from the parental BAC containing the SARS-CoV-2 USA-WA1/2020 strain genome described in Ye et al. (2020); the 15.4 kb fragment was generated by restriction digestion. For mutagenesis, the N gene-containing fragment was further broken up into three subfragments; amino acid and TRS mutations were introduced with overlapping forward and reverse mismatch primers and invariant end primers (Table 1), and the full-length mutant fragments were then assembled and amplified by overlap extension PCR. KR^+TRS^ mutation was modeled after the SARS-CoV-2 Gamma variant.

**Table 1.**
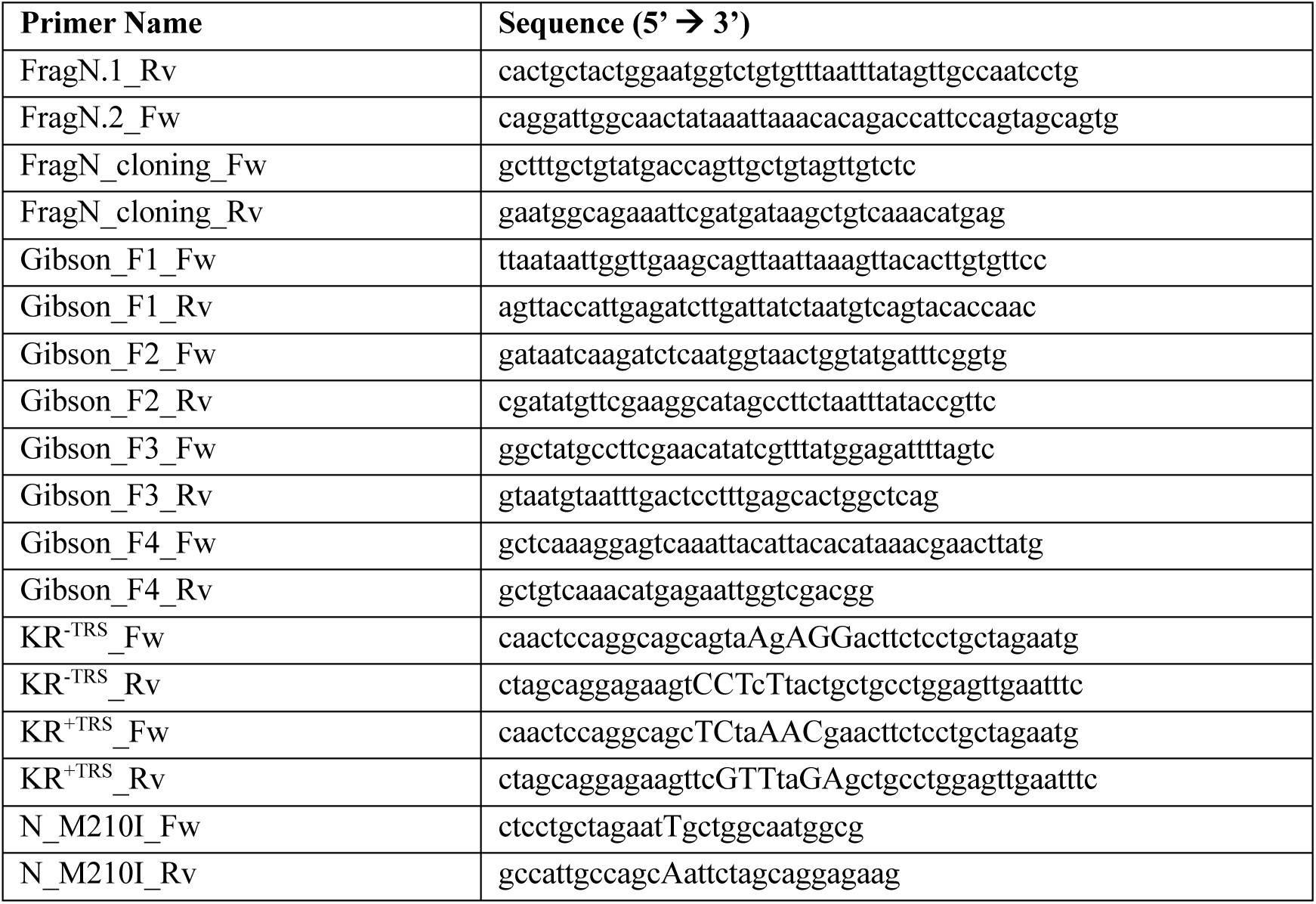
Amplification and mutagenesis primers (mismatches in uppercase)

All Gibson assembly fragments were gel-extracted and assembled using the Gibson Assembly Master Mix (New England Biolabs). Fragments were combined at equimolar ratios (15 fmol each) in 30 µL reactions that were incubated at 50℃ for 60 minutes according to the manufacturer’s instructions. All mutant BACs were validated by restriction digest and Sanger sequencing of the entire viral genome using 56 sequencing primers (Table 2). Verified BAC clone DNA was purified for transfection using plasmid Midi Kit (Qiagen). PCR cycling conditions for amplification of the Gibson fragments from the parental BAC: *95℃-1 min; 30 × (95℃ - 15 sec; 60℃ - 30 sec; 72℃ - 5 min); 72℃ - 5 min.* PCR cycling conditions for mutagenesis and sequential overlap extension reactions: *95℃-1 min; 15-25 × (95℃ - 15 sec; 60-63℃ - 30 sec; 72℃ - 2-5 min); 72℃ - 5 min*.

**Table 2.**
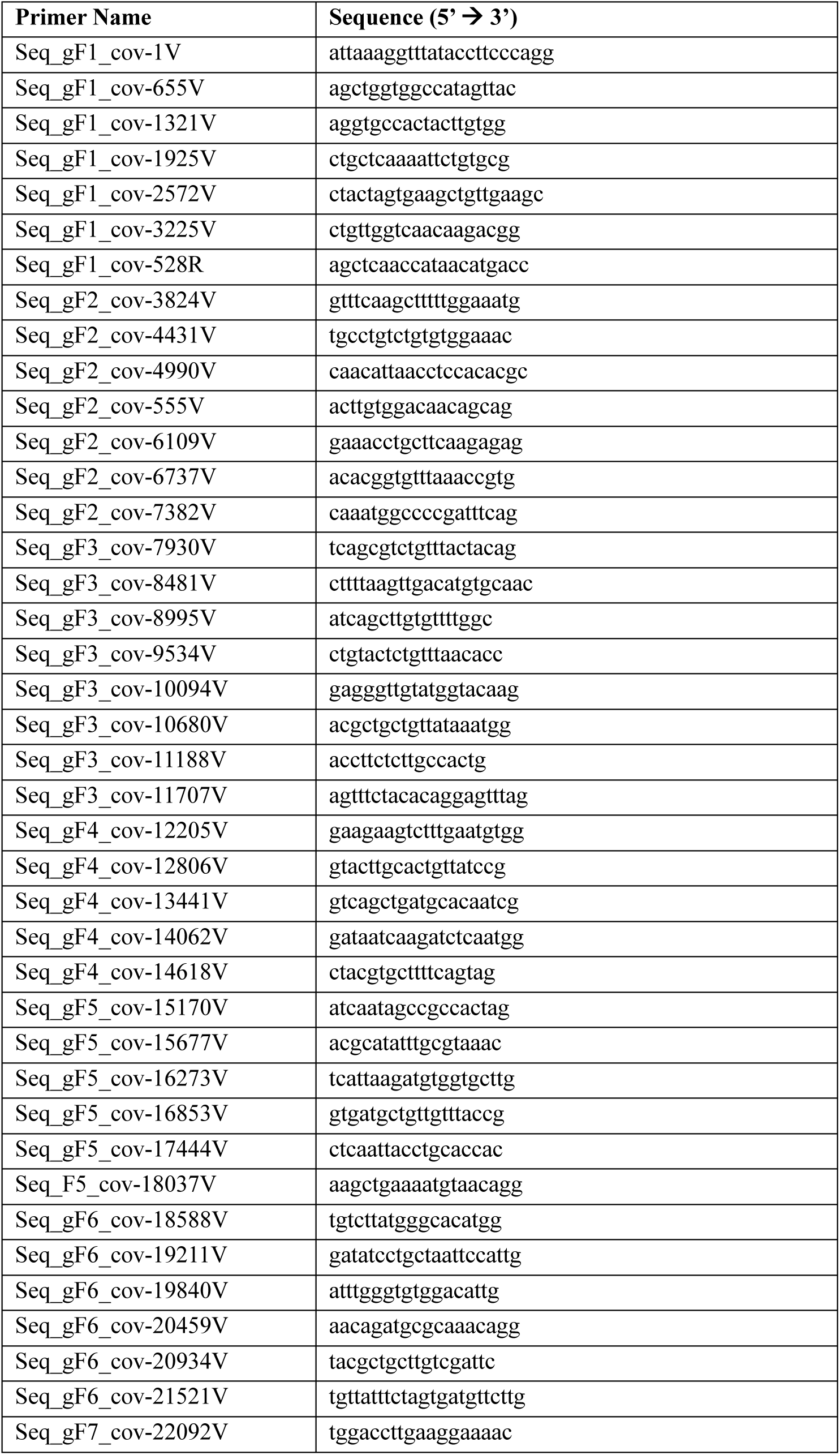
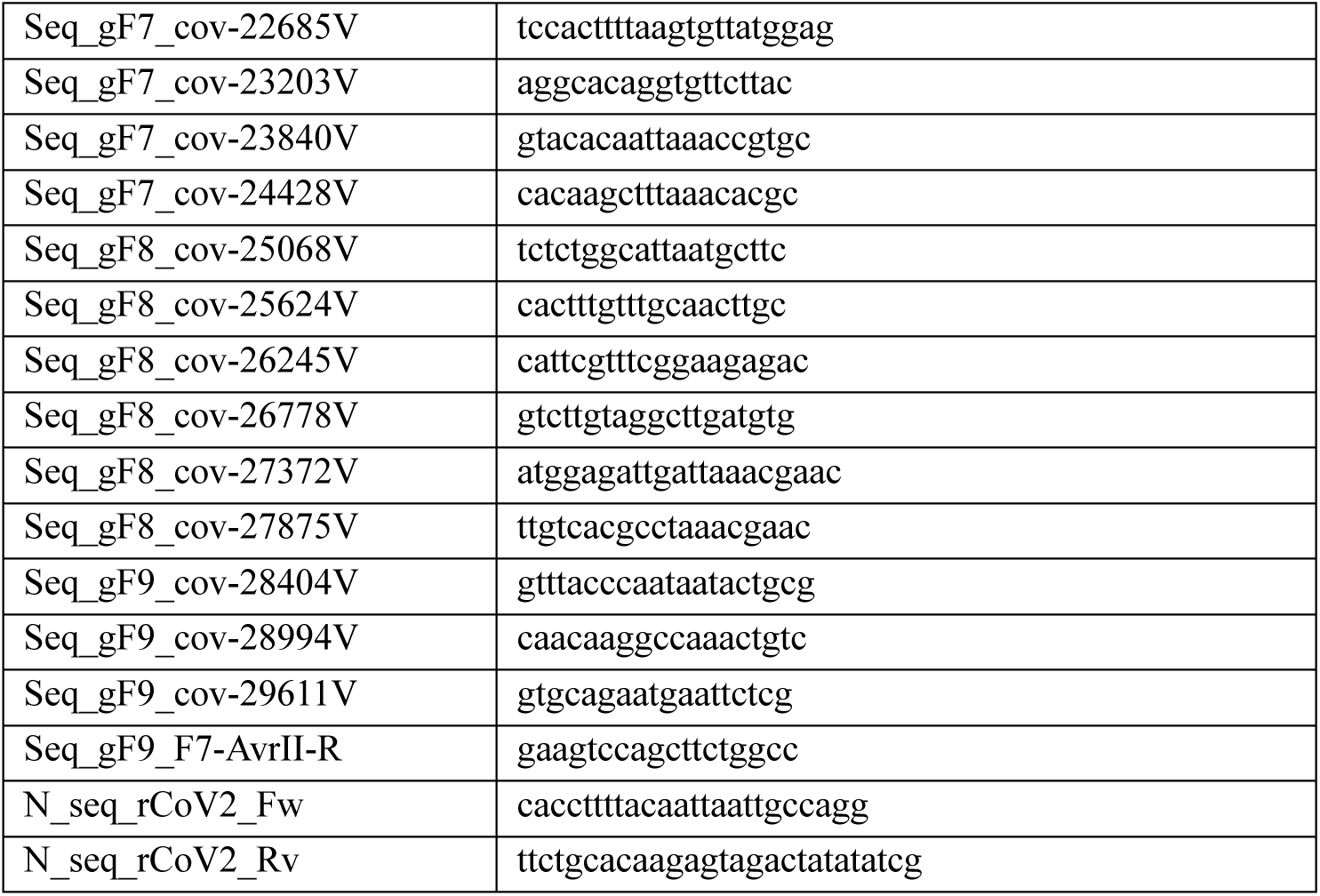
rSARS2-BACmid sequencing primers.

### Recombinant virus generation and validation

Recombinant SARS-CoV-2 were made after consultation with the Public Health Agency of Canada (PHAC), the University of Calgary’s Biosafety Office, and the Containment Level 3 (CL3) Oversight Committee. VeroE6^TMPRSS2^ cells were seeded into 6-well plate. 24 hours after seeding, 0.5 µg of recombinant SARS-CoV-2 BAC was combined with 1.5 µL of Lipofectamine 2000 (Thermo-Fisher). Transfection mixture was added to cells containing antibiotic-free DMEM. Once cytopathic effect was observed (48-96 hpi), virus-containing supernatant was harvested, cell debris was removed by centrifugation, and virus was frozen at -80°C for single use. Viral RNA was harvested from passage 0 and passage 1 stocks using QIAamp Viral RNA Mini kit (Qiagen) as per manufacturer’s instructions, without including carrier RNA. Recombinant viruses were propagated in VeroE6^TMPRSS2^ cells. All recombinant virus experiments were conducted with passage 1 virus stocks.

### Next generation sequencing of recombinant viruses

Viral RNA was prepared for whole genome sequencing via multiplexed PCR amplicon tiling using the SARS-CoV-2 Midnight Amplicon Panel (N. E. Freed et al., 2021) (IDT; this study used Midnight Amplicon Panel v1). A detailed protocol is available and was followed up to and including step 11 (N. Freed & Silander, 2021). Briefly, cDNA was prepared using LunaScript RT Supermix Kit (NEB) according to the manufacturer’s specifications, with 8 μL of viral RNA as template. cDNA was then used in two PCR reactions with the Midnight Amplicon Panel to generate two unique amplicon pools which were then mixed at a 1:1 ratio. Combined amplicon pools were submitted to the University of Calgary Centre for Health Genomics and Informatics for library preparation followed by sequencing via Illumina MiSeq 300-cycle nano (2 x 150 reads). Resulting raw single-end sequence Illumina reads were pre-processed using fastp (S. Chen et al., 2018), under default parameters, for the trimming of Illumina adaptors and low-quality reads; trimmed reads were exported as .fastq.gz files. Trimmed reads were then imported to Geneious Prime (v.2025.0.1) to generate paired end (inward pointing) reads with insert size=“150”. Paired reads were further processed using two Genious Prime-resident processing tools: Dedupe (v.38.84 by Brian Bushnell) for the removal of duplicate reads, followed by BBNorm (v.38.84 by Brian Bushnell) (Haider et al., 2014) for paired read error correction and normalization. Dedupe was run under default parameters, and BBNorm was run under default parameters for error correction and under the following parameters for normalization: Target Coverage Level=“220”, Minimum Depth=“6”. Fully processed paired reads were then mapped against the SARS-CoV-2 reference genome (NC_045512) which was modified to replace the nucleotides in positions 28,877 – 28,885 with nucleotide wild cards (“N”). This permits an unbiased sequence alignment albeit the mutational differences in this wild card region between wild-type and mutant viral sequences. Reference mapping parameters: Sensitivity=”Medium Sensitivity/Fast”, Minimum mapping quality=“30”. The Find Variants/SNPs tool was employed with Minimum Variant Frequency=“0” for detection of all possible sub-populations under the reference sequence wild cards. The wild card region had a coverage of ∼330 across sequenced viral species. Consensus sequences were exported with a consensus threshold set to 75%. The consensus sequence for wild-type virus matched the reference genome with ≥ 99.4% likeness across the wild card region; the consensus sequence for mutant viruses matched the expected sequences based on the intended mutations made with mutational variant frequencies ≥ 97.2% across the wild card region, for all recombinant viruses.

### TaqMan RT-qPCR analysis of viral co-infection competition assays

Determination of transcript viral copy number in viral co-infection competition assays was conducted via a probe-based RT-qPCR TaqMan assay to differentiate viral species. Primer and TaqMan probe sequences can be found in Table 3; all probes contain a 5’ fluorescent reporter and a 3’ non-fluorescent quencher (NFQ) and minor groove binding (MGB) protein. Qubit 1× dsDNA High Sensitivity Assay Kit (Invitrogen) was used to accurately determine competition assay standard plasmid concentrations with a Qubit 3.0 Fluorometer (Invitrogen). The plasmid copy number of standards was calculated and used to prepare a 6-point standard curve between 1E2 to 1E7 known plasmid copies under a 1:10 dilution series. TaqMan probes were validated against their respective standard curve target; reaction efficiencies for all probes were between 90 – 95% with R^2^ values > 0.9990. Probes were also confirmed to be specific for their respective standard target, displaying no cross-reactivity when tested against off-target standards. For standard curve analysis, RT-qPCR reactions were prepared in duplicate per standard dilution. For co-infection competition assay samples, RT-qPCR reactions were prepared in quadruplicate per sample, divided into a duplicate for detection of one viral species and a duplicate for detection of the other viral species. Reaction consisted of (final concentrations) 500 nM forward and reverse SARS-CoV-2 N TaqMan assay primers, 250 nM of respective probe, 5 μL 2× TaqMan Universal PCR Master Mix (Thermo-Fisher) containing uracil-N-glycosylase (UNG) for the prevention of carryover contamination, and either 2.5 μL of plasmid standards at 0.4E2 – 0.4E7 standard plasmid copies/μL (final of 1E2 – 1E7 plasmid copies in reaction) or 2.5 μL of diluted cDNA sample. In general, cDNA was diluted 1:50 for HUVEC-originating samples, 1:100 for A549-originating samples, and 1:500 for HEK293A-ΔG3BP1- and VeroE6-originating samples. Reaction cycling conditions were as follows: *UNG activation at 50°C for 2 min, initial denaturation at 95°C for 10 min, cycling at 40× (denaturation at 95°C - 15 s, annealing and amplification at 60°C - 30 s.* Resulting Cq values were used to determine the number of viral transcript copies of a single viral species detected in the reaction by relating it against the corresponding standard curve. The total number of viral transcripts detected in the reaction was determined by taking the sum of detected viral transcript copies across both species. The percentage population of individual viral species within a reaction was expressed as a relative ratio to the total viral transcript copy number detected. TaqMan probe specificity was validated in RT-qPCR assays using cDNA from RNA isolated from recombinant SARS-CoV-2-infected HUVEC^ACE2^ cells (Fig S2D).

**Table 3.**
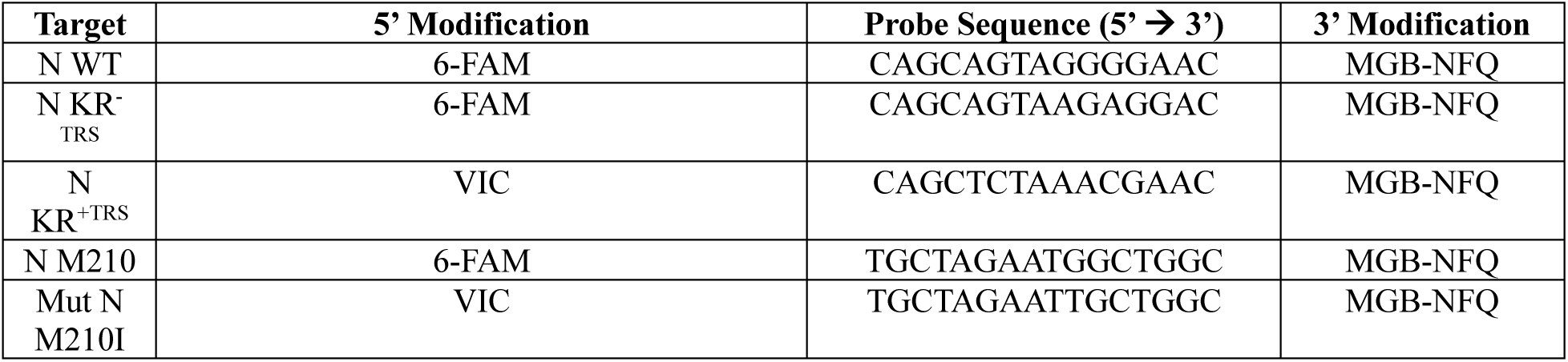
TaqMan primer sequences.

### Virus propagation, titration, and infection

Experiments with SARS-CoV-2 variants and recombinant SARS-CoV-2 viruses were conducted in a CL3 facility, and all standard operating procedures were approved by the CL3 Operations Oversight Committee and Biosafety Office at the University of Calgary. SARS-CoV-2 Toronto-01 (TO-1), Alpha, Beta, Gamma, and Delta isolates were titrated and propagated in VeroE6 cells, as in Kleer, et al. 2022. Omicron BA-1 isolate (a generous gift from Dr. Lorne Tyrrell, Dr. Michael Joyce, and Holly Bandi [University of Alberta]) and all recombinant SARS-CoV-2 viruses were titrated and propagated in VeroE6^TMPRSS2^ cells. Virus infectious titer was enumerated by plaque assay on VeroE6 or VeroE6^TMPRSS2^ (Omicron and rSARS-2) cells using equal parts 2.4% w/v semi-solid colloidal cellulose overlay (Sigma; prepared in ddH_2_O) and 2X DMEM (Wisent) with 1% FBS, 100 U/mL penicillin, 100 µg/mL streptomycin, 2 mM L-glutamine. 48-72 hours post infection cells were fixed with 4% formaldehyde and stained with 1% w/v crystal violet (Thermo-Fisher) for plaque enumeration. For propagation, cells were infected at an MOI of 0.01 for 1 hour in serum-free, antibiotic-free DMEM at 37°C. After virus adsorption, media was replaced with fresh media, containing 2% FBS and 100 U/mL penicillin, 100 µg/mL streptomycin, 2 mM L-glutamine. 3-5 days post infection, virus-containing supernatant was harvested and cell debris was removed by centrifugation. Virus stocks were stored at -80°C and infectious titer was determined by plaque assay.

For experiments, cells were seeded to achieve ∼80% confluence at the time of infection. Virus inoculum was diluted in serum-free, antibiotic-free DMEM to achieve the desired MOI. Growth media was removed from cells and replaced with the diluted virus inoculum. Cells were infected for 1 hour at 37°C, after which the media was replaced with complete growth media until experimental endpoint.

### Extracellular virus particle concentration

VeroE6 cells were infected with SARS-CoV-2 TO-1 or Alpha (MOI=1) in T75 cm^2^ flasks. 24 hpi, virus-containing supernatant was collected, and cell debris was removed by low-speed centrifugation (1500 RPM for 5 mins) and filter sterilization using a 0.45µm filter (VWR). Mature viral particles were concentrated as in (Bo et al., 2020). Briefly, viruses were concentrated by high-speed centrifugation (20,000 RPM for 4 hours; Beckman Coulter Optima LK-90, with T70.1 fixed-angle rotor) through a 20% (w/v in PBS) sucrose cushion. Viral pellet was resuspended in 2X Laemmli buffer and boiled at 92°C for 15 minutes.

### Fluorescent RNA oligonucleotides

Complementary RNA oligonucleotides with the following sequences were designed as in (Lodola et al., 2023) and purchased from Thermo-Fisher:

**38 nt:** 5’-AUGAAGGUUUGAGUUGAGUGGAGAUAGUGGAGGGUAGU-3’

**Fluorescein-18 nt:** 5’-fluorescein-CUCAACUCAAACCUUCAU-3’ Oligonucleotides were resuspended in Nuclease-Free Duplex Buffer (30 mM HEPES, pH 7.5; 100 mM potassium acetate) (Integrated DNA Technologies). To obtain dsRNA, 38 nt and fluorescein-18 nt oligonucleotides were combined at an equimolar ratio for a final concentration of 100 µM, heated for 5 minutes at 95°C, then gradually cooled to 4°C. RNA oligonucleotide transfections were performed using Lipofectamine 2000 (see Methods “Transient transfections and drug treatments”).

### Reverse transcriptase quantitative and non-quantitative-PCR

Intracellular RNA was harvested using RNeasy Mini kit (Qiagen) according to manufacturer’s instructions. Extracellular RNA was extracted using QIAamp Viral RNA Mini kit (Qiagen) as per manufacturer’s instructions, without including carrier RNA. Extracted RNA was stored at -80°C until use. RNA concentrations were determined using NanoDrop One^C^ (Thermo-Fisher) and 1000 ng of RNA was converted to cDNA using Maxima H Minus Reverse Transcriptase (Thermo-Fisher) according to manufacturer’s instructions. For RT-PCR, following PCR amplification with Taq DNA polymerase (Thermo-Fisher), amplicons were resolved by agarose gel electrophoresis on a 1.5% gel with Syber Safe (Thermo-Fisher). RT-PCR cycling conditions: initial denaturation at *95℃ - 30 sec; cycling at 30* x *(95℃ - 30 sec; annealing at 59℃ - 30 sec; extension at 68℃ - 5 sec); final extension at 68℃ - 5 min.* Resolved amplicons were imaged using the ChemiDoc MP Imaging System. For RT-qPCR, cDNA was diluted 1:10 with nuclease-free H_2_O and cDNA was amplified using SsoFast EvaGreen Master Mix (BioRad). RT-qPCR cycling conditions: initial denaturation at *98℃ - 2:00 min; cycling at 39* x *(98℃ - 0.02 min; annealing and amplification at 60℃ - 0.05 min); final extension at 65℃ - 0.10 min, followed by melt curve analysis (60℃ - 0.10 min and 95℃ - 0.2 min)*. Relative RNA abundance was calculated using the 2^^-ΔΔCt^ equation, with 18S rRNA as a housekeeping control gene. Primer sequences can be found in Table 4.

**Table 4.**
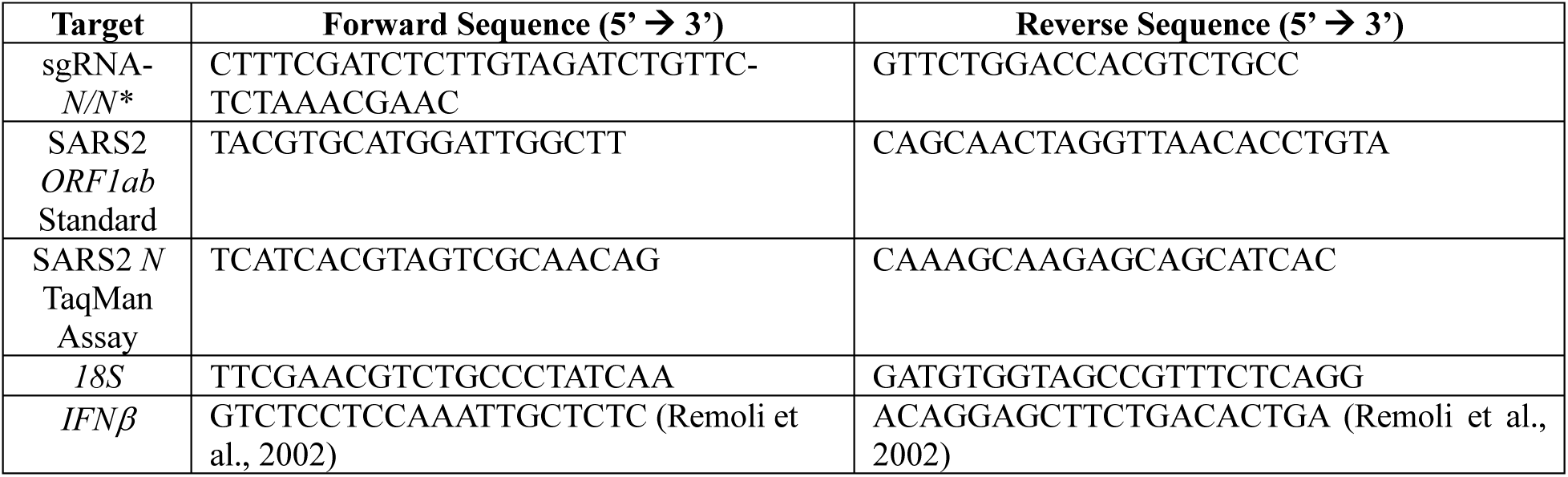
RT-PCR and qPCR Primer sequences.

### Transient transfections and drug treatments

Transient transfections in HeLas were conducted using Fugene HD (Promega) according to the manufacturer’s guidelines. Briefly, in Opti-MEM (Gibco), plasmid DNA and Fugene HD were added, with 1µg of DNA to 3µL of Fugene HD. The transfection mixture was added dropwise to cells containing antibiotic-free media. 24 hours post transfection, the media was replaced. Poly I:C and poly A:U (Invitrogen) transfections were conducted with Fugene HD, as described above. RNA oligonucleotide transfections were conducted with Lipofectamine 2000 (Thermo-Fisher) according to the manufacturer’s guidelines. In Opti-MEM, RNA oligonucleotides and Lipofectamine 2000 were added in a 1:1 ratio (1 µg of RNA to 1 µL of Lipofectamine 2000). Transfection mixture was added dropwise to cells containing antibiotic-free media. All other transfections were achieved using polyethylimine (PEI, Polysciences). In Opti-MEM, plasmid DNA and PEI (1mg/mL) were added, with 1µg of DNA to 3µL of PEI. Transfection mixture was added dropwise to cells containing serum-free, antibiotic-free media. 6 hours post transfection, the media was replaced. Sodium arsenite (SA; Sigma) was added to cells at a 500 µM final concentration.

### Production and use of recombinant lentiviruses

Second generation recombinant lentiviruses were produced as in Kleer et al. (2022). Briefly, HEK293T cells were transfected with pMD2.G, psPAX2, and the lentiviral transfer plasmid at a ratio of 1:2:3.3 (µg) using PEI as described above. 6 hours post transfection, the media was replaced with DMEM containing 10% serum, but no antibiotics. 48 hours post transfection, virus supernatant was harvested, filtered with a 0.45µm syringe filter (VWR), and stored at -80°C for single use. For transductions, lentiviruses were thawed at 37°C and added to target cells in complete media containing 5 µg/mL polybrene (Sigma). 24 hours post transduction, media was replaced with complete media containing puromycin (1 µg/mL; Thermo-Fisher) or blasticidin (5 µg/mL; Thermo-Fisher). Cells were selected for 48 hours before replacing with complete media or until experimental endpoint.

### Co-immunoprecipitations

Co-immunoprecipitations were conducted as in (Robinson et al., 2023). Briefly, HEK293T cells were seeded into a 6-well plate. Co-immunoprecipitations were conducted 48 hours post plasmid transfection. Cells were transfected with poly I:C or mock transfected 3 hours before conducting co-immunoprecipitation. Cells were resuspended in lysis buffer (150 mM NaCl, 10 mM Tris pH 7.4, 1 mM EDTA, 1% v/v Triton X-100, 0.5% v/v NP-40, Roche protease inhibitor tablet). For Fig 4A, 10 or 50 µg/mL of RNase A (Thermo-Fisher) was added to lysis buffer and incubated for 30 minutes at room temperature. Following lysis, cells were incubated with the indicated primary antibody overnight at 4°C. Protein G magnetic Dynabeads (Thermo-Fisher) were incubated in blocking buffer (150 mM NaCl, 10 mM Tris pH 7.4, 1 mM EDTA, 5 mg/mL BSA) overnight at 4°C. Immunoprecipitation was conducted and beads were boiled in 4X Laemmli buffer (BioRad) with 10% v/v β-mercaptoethanol.

### Immunoblotting and densitometric analysis

Cells were lysed in 2X Laemmli buffer and stored at -20 until use. For SARS-CoV-2-infected cells, cells were lysed in 2X Laemmli buffer and boiled at 92°C for 15 minutes, then stored at -20°C. Protein concentration was quantified for normalization using DC Protein Assay kit (BioRad) according to manufacturer’s instructions. 6-15 µg of protein lysate was resolved by SDS-PAGE using TGX Stain-Free acrylamide gels (BioRad). After protein transfer, total protein images were acquired from PVDF membranes on the ChemiDoc MP Imaging System (BioRad). PVDF membranes were blocked with 5% BSA (Thermo-Fisher) in TBST (Tris-buffered saline solution with 0.1% Tween-20). Primary antibodies and secondary antibodies were diluted in 5% BSA in TBST and 5% skim milk in TBST, respectively, except when probing for pIRF3, where the secondary antibody was diluted in 5% BSA. Antibody concentrations can be found in Table 5. Densitometry was conducted in ImageLab software (BioRad) by normalizing to total protein.

**Table 5.** Antibodies.

| Antibody | Species | Vendor/Catalog # | Application | Dilution |
| --- | --- | --- | --- | --- |
| ACE2 | Rabbit | CST/4355S | Immunoblot | 1:1000 |
| Actin-HRP conjugated | Rabbit | CST/5125S | Immunoblot | 1:10 000 |
| FLAG | Chicken | Abcam/ab1170 | Immunofluorescence | 1:100 |
| FLAG | Mouse | CST/8146S | Immunoblot<br>Immunoprecipitation | 1:1000<br>1:1000 |
| FLAG | Rabbit | CST/14793T | Immunoblot<br>Immunofluorescence | 1:1000<br>1:1000 |
| G3BP1 | Mouse | BD/611126 | Immunoblot<br>Immunofluorescence | 1:1000<br>1:1000 |
| Hedls | Mouse | Santa Cruz/8418 | Immunofluorescence | 1:1000 |
| HA | Rabbit | CST/3724S | Immunofluorescence | 1:1000 |
| IRF3 | Rabbit | CST/11904T | Immunoblot | 1:1000 |
| pIRF3 | Rabbit | CST/37829 | Immunoblot | 1:500 |
| J2 | Mouse | Millipore /MABE1134 | Immunofluorescence | 1:100 |
| J2 (the above was discontinued) | Mouse | CST/76651L | Immunofluorescence | 1:500 |
| PKR | Rabbit | Abcam/ab184257 | Immunoblot | 1:1000 |
| pPKR | Rabbit | Abcam/ab32036 | Immunoblot | 1:1000 |
| SARS-CoV-2 N | Rabbit | Novus/NBP3-05730 | Immunoblot<br>Immunofluorescence | 1:1000<br>1:1000 |
| TIAR | Rabbit | CST/8509T | Immunofluorescence | 1:800 |
| Alexa Fluor 405 anti-rabbit IgG | Donkey | Abcam/ab175651 | Immunofluorescence | 1:200 |
| Alexa Fluor 488 anti-rabbit IgG | Chicken | Invitrogen (A21441) | Immunofluorescence | 1:1000 |
| Alexa Fluor 488 anti-mouse IgG | Goat | Invitrogen (A11029) | Immunofluorescence | 1:1000 |
| Alexa Fluor 555 anti-mouse IgG | Donkey | Invitrogen (A31570) | Immunofluorescence | 1:1000 |
| Alexa Fluor 647 anti-mouse IgG | Chicken | Invitrogen (A21463) | Immunofluorescence | 1:1000 |
| Alexa Fluor 488 anti-chicken IgY | Goat | Invitrogen (A11039) | Immunofluorescence | 1:200 |

### Poly I:C-biotin pulldowns

The poly I:C-biotin pulldown protocol was adapted from (Schoen et al., 2023). HEK293T cells were seeded into a 6-well plate. Lysates were collected 24 hours post plasmid transfection. The media was removed and cells were washed with ice-cold 1X PBS, then harvested in ice-cold 1X PBS and pelleted by centrifugation. Cells were then resuspended in ice-cold 1X PBS, pelleted by centrifugation, and resuspended in lysis buffer (150 mM NaCl, 10 mM Tris pH 7.4, 1 mM EDTA, 1% v/v Triton X-100, 0.5% v/v NP-40, 0.1% v/v P8340 protease inhibitor cocktail (Sigma Aldrich)). Cell lysates were combined with 1 U/μL RNase T1 (Thermo-Fisher) and rocked for 30 minutes at room temperature, then rotated at 4°C for 30 minutes and subjected to centrifugation to remove cell debris.

M270 Streptavidin magnetic Dynabeads (Thermo-Fisher) were washed three times with 1X binding and washing (B&W) buffer (2 M NaCl, 1 mM EDTA, 10 mM Tris-HCl pH 7.4). Beads were combined with 1X B&W buffer containing 300 ng poly I:C (HMW)-biotin (Invivogen) and 200 U/mL RNaseOUT Recombinant Ribonuclease Inhibitor (Thermo-Fisher) and rotated at 4°C for 2.5 hours, then washed three times with 1X B&W buffer containing 200 U/mL RNaseOUT Recombinant Ribonuclease Inhibitor. Cell lysates were combined with beads and rotated at 4°C overnight. Beads were washed three times with wash buffer (150 mM NaCl, 10 mM Tris pH 7.4, 1 mM EDTA), then elution was performed by boiling beads in 4X Laemmli buffer (BioRad) with 10% v/v β-mercaptoethanol.

### rRNA integrity assay

Cells were transfected with 1.0 µg high molecular weight poly I:C (Invitrogen) for 3 hours. Intracellular RNA was harvested using RNeasy Mini kit (Qiagen) according to manufacturer’s instructions. Extracted RNA was stored at -80°C until use. RNA concentrations were determined using NanoDrop One^C^ (Thermo-Fisher). Extracted RNA was diluted to a final concentration of 50 ng/µL, then submitted to the University of Calgary Centre for Health Genomics and Informatics for automated electrophoresis using an Agilent 4200 TapeStation system.

### Immunofluorescence

Cells were seeded onto 18mm glass round-bottom #1.5 coverslips (Electron Microscopy Sciences). At experimental endpoint, cells were fixed with 4% paraformaldehyde (Electron Microscopy Sciences) in PBS (Gibco) for 10 minutes at room temperature, or 30 minutes if infected with SARS-CoV-2. Cells were permeabilized with 0.1% (v/v) Triton X-100 (Sigma-Aldrich) in 1X PBS for 10 minutes at room temperature. Cells were blocked with 1% (v/v) human AB serum (Sigma-Aldrich) in PBS for 60 minutes at room temperature. Primary and secondary antibodies were diluted in 1% human AB blocking buffer at listed concentration (Table 5). Primary antibodies were incubated over night at 4°C, secondary antibodies were incubated for 60 minutes at room temperature. Nuclei were stained with 1µg/mL Hoechst (Invitrogen). Coverslips were mounted with antifade Prolong Gold (Thermo-Fisher). Unless otherwise stated, cells were co-stained with Alexa Fluor 647 and Alexa Fluor 488 to minimize fluorophore cross excitation and bleed through. Images were captured using a Zeiss AxioObserver Z1 microscope with a 40X oil-immersion objective for all images used for SG and P-body quantification. For co-localization analysis and all representative images, images were acquired using a Zeiss LSM 880 confocal microscope using a 63X oil-immersion objective. Exposure time and laser power was kept consistent within replicates.

### RNP granule quantification and enrichment analysis

P-bodies and SGs were quantified using CellProfiler4.0.6 (cellprofiler.org), as in Kleer et al. (2022). Nuclear staining was used to identify individual cells by applying manual thresholding followed by “Identify Primary Objects”. Acceptable nuclei size was cell-type specific and dependent on imaging objective. Cell boarders were defined by applying a “Propagation” function from each nucleus. Propagation distance was dependent on imaging objective and was cell-type specific to account for differences in cell size. Nuclei in close proximity have automatically reduced propagation distance, such that each cytoplasm is mutually exclusive and therefore cytoplasmic area or granules are not double counted. As P-bodies and SGs are cytoplasmic, the nuclear area was subtracted using “Mask” and “Remove” functions. The cytoplasmic area was masked and RNP granules were measured within each cell. Cells were now stratified into ‘Positive Cells’ and ‘All Cells’. Positive Cells are those that stain positive for N (virus infection) or FLAG (over-expression). ‘All Cells’ include cells expressing N/FLAG and non-expressing cells. In this way, in control treatments (i.e. mock-infected cells or EV-transfected cells), RNP granules in All Cells are calculated. Whereas, in experimental treatments (i.e. infected cells or N-transfected cells), RNP granules in only Positive Cells are calculated. To identify RNP granules, granules were thresholded to reduce background staining. Granules were defined based on size; P-bodies were defined as Hedls-positive puncta ranging from 3-13 pixels and SGs were defined as G3BP1-positive puncta ranging from 2-50 pixels for poly I:C-induced SGs and 3-50 pixels for SA-induced SGs. Quantification parameters were identical within independent experiments; however, image thresholding was altered between independent experiments to account for staining variability. P-bodies and SGs were quantified and graphed as number of foci per cell.

For enrichment of N proteoforms in puncta dsRNA foci, images were all acquired on Zeiss LSM 880 confocal microscope with z-stacks. Nuclei and cell boarders were defined as above, however as dsRNA foci can be found overlapping with nuclei, the nucleus was not subtracted from the total cellular area. Nucleocapsid staining was thresholded and N positive cells were identified as above. dsRNA foci were identified by thresholding and size exclusion as above. To measure nucleocapsid localization inside dsRNA foci, puncta were subjected to a “Mask” and “Keep” function. To measure nucleocapsid localization outside of SGs or dsRNA foci, puncta were subjected to a “Mask” and “Remove” function. The mean integrated intensity of unthresholded N staining from “Nucleocapsid Inside Puncta” and “Nucleocapsid Outside Puncta” were measured. Percent enrichment of N in puncta was calculated by addition of the mean integrated intensity of N Inside Puncta divided by the total cellular integrated intensity (integrated intensity of ‘Nucleocapsid Inside Puncta’ plus integrated intensity of ‘Nucleocapsid Outside Puncta’). N-puncta enrichment was calculated and represented per cell. Quantification parameters were identical within independent experiments; however, image thresholding was altered between independent experiments to account for staining variability.

### Statistics

All statistical analyses were performed using GraphPad Prism 9.0. RNP granule (SGs and P-bodies) per cell foci counts as well as per cell enrichment quantifications were plotted such that independent biological replicates were combined and plotted on one graph. These per cell data are naturally skewed and therefore non-parametric (as determined by the Shapiro-Wilk normality test), therefore we chose used rank-sum statistical analyses (Mann-Whitney U test for comparing two independent groups, and Kruskal Wallis test for comparing multiple independent groups). All other statistics were performed using parametric statistical analyses (T-test for comparing two independent groups or ANOVA for comparing multiple independent groups) unless otherwise indicated.

## Supporting information

All supplemental figures

## ACKNOWLEDGEMENTS

We sincerely thank Dr. Denys Khaperskyy (Dalhousie University) for providing HEK293T-ΔG3BP1 cells and for helpful discussions. We would like to thank Dr. Roy Duncan (Dalhousie University), Dr. Craig McCormick (Dalhousie University), and Dr. James Burke (University of Florida Scripps Institute) for helpful discussions about this work and Dr. Luis Martinez-Sobrido and Dr. Chengjin Ye (Texas Biomedical Research Institute) for the SARS-CoV-2 BACmid system and helpful discussions regarding recombinant SARS-CoV-2 cloning. This work would not have been possible without Drs. Anne Vaahtokari and Luc Provencher of the Charbonneau Microscopy Facility and Drs. Devender Kumar and Shaunna Huston of the University of Calgary for CL3 support. Finally, we would like to thank all the members of the Corcoran lab for helpful discussions about experimental design. RPM was supported by a Snyder Institute Beverley Phillips Doctoral training award and a CIHR CGS-D award. MPBM was supported by a CSM graduate training award, a Snyder Institute Beverley Phillips Doctoral training award and an Alberta graduate scholarship. NS was supported by a CIHR CGS-M scholarship and a CSM graduate training award. This study was supported by operating funds awarded to JAC from the Canadian Institutes for Health Research (CIHR) (project grant #195645) and an operating grant (#175622) to the Coronavirus Variants Rapid Response Network (CoVaRR-Net), of which JAC is a member.

## AUTHOR CONTRIBUTIONS

**Rory P Mulloy:** Conceptualization; Data curation; Formal analysis; Investigation; Methodology; Validation; Visualization; Writing – original draft; Writing – review & editing

**Danyel Evseev:** Data curation; Investigation; Methodology

**Maxwell P. Bui-Marinos:** Data curation; Formal analysis; Investigation; Methodology

**Noga Sharlin:** Data curation; Investigation; Methodology

**Jennifer A. Corcoran:** Conceptualization; Funding acquisition; Project administration; Resources; Supervision; Writing – original draft; Writing – review & editing

### Competing interests

The authors have declared that no competing interest exist.

## Notes

### Competing Interest Statement

The authors have declared no competing interest.

