## Supplementary material for "A truncated SARS-CoV-2 nucleocapsid protein enhances virus fitness by evading antiviral responses": All supplemental figures

Figure S1

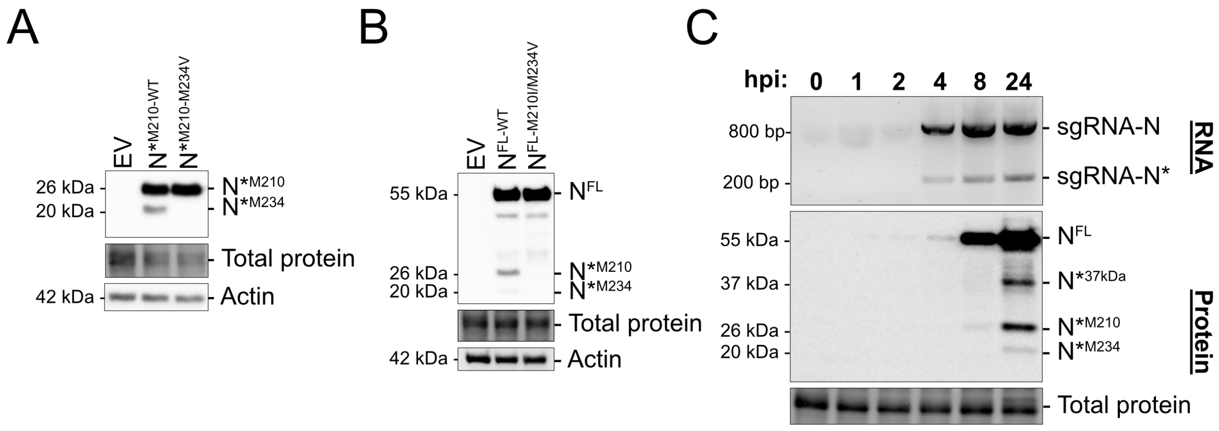

A
EV
N*M210-WT
N*M210-M234V
26 kDa
20 kDa
N*M210
N*M234
Total protein
42 kDa
Actin
B
EV
NFL-WT
NFL-M210I/M234V
55 kDa
NFL
26 kDa
20 kDa
N*M210
N*M234
Total protein
42 kDa
Actin
C
hpi: 0 1 2 4 8 24
800 bp
200 bp
sgRNA-N
sgRNA-N*
RNA
55 kDa
37 kDa
26 kDa
20 kDa
NFL
N*37kDa
N*M210
N*M234
Protein
Total protein

**Figure S1. Methionine 210 and 234 enable truncated N production.**

**A.** $N*^{M210\text{-}WT}$, or $N*^{M210\text{-}M234V}$ containing a C-terminal FLAG tag, or an empty vector (EV) control were over-expressed in HEK293T cells and protein lysate was subjected to SDS-PAGE and immunoblotting with anti-FLAG and anti-actin antibodies.

**B.** $N*^{M210\text{-}WT}$, or $N^{FL\text{-}M210I/M234V}$ containing a C-terminal FLAG tag, or an empty vector (EV) control were over-expressed in HEK293T cells and protein lysate was subjected to SDS-PAGE and immunoblotting with anti-FLAG and anti-actin antibodies.

**C.** Kinetics of N proteoform and sgRNA profile was determined by infecting Calu3 cells with SARS-CoV-2 Alpha variant (MOI=4). At the indicated time post-infection, RNA and protein lysate was harvested and was subjected to RT-PCR and agarose gel electrophoresis, and SDS-PAGE and immunoblotting, respectively.

Figure S2

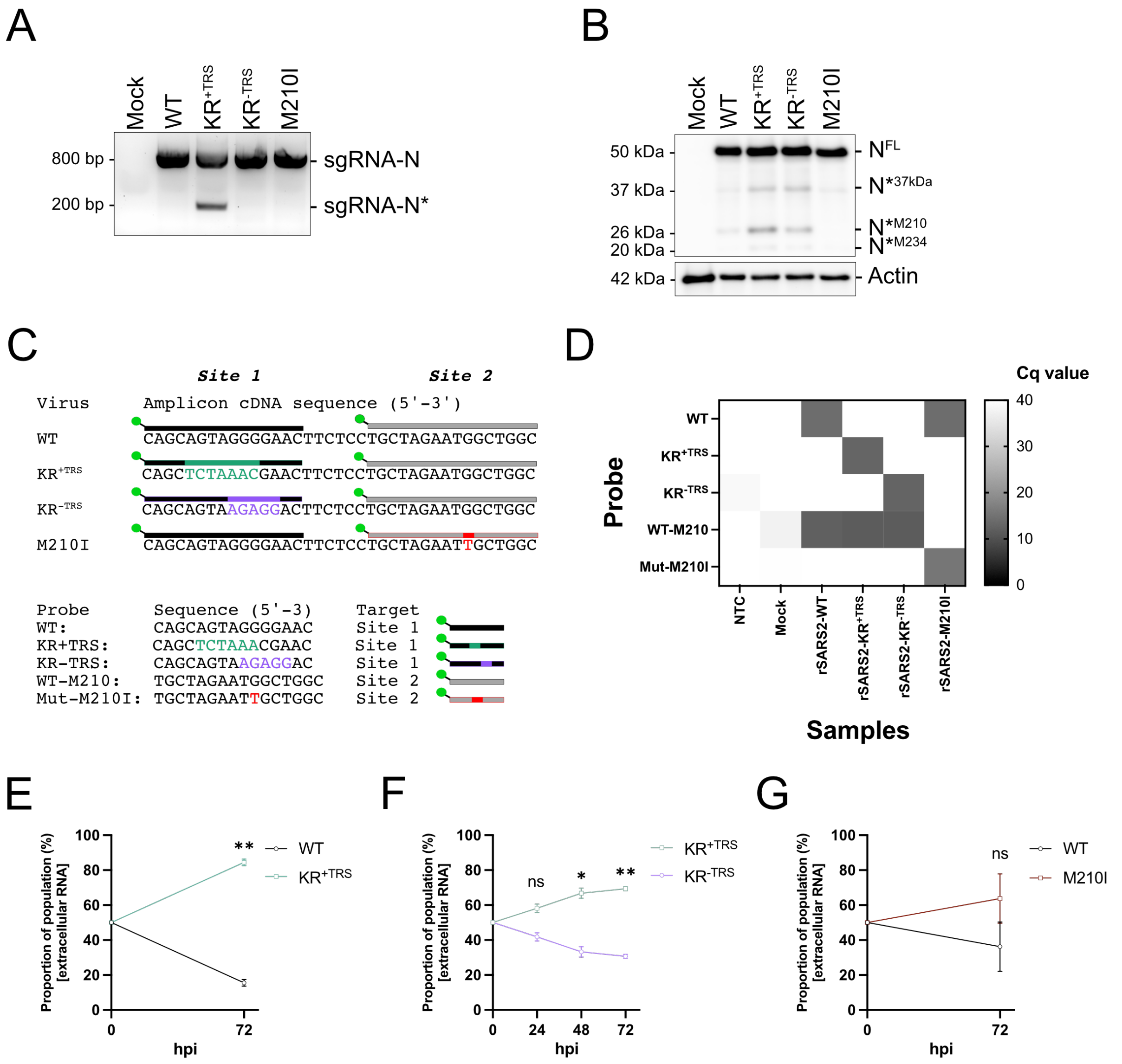
A
Mock
WT
KR+TRS
KR-TRS
M210I
800 bp
200 bp
sgRNA-N
sgRNA-N*
B
50 kDa
37 kDa
26 kDa
20 kDa
42 kDa
NFL
N*37kDa
N*M210
N*M234
Actin
C
Site 1
Site 2
Virus
Amplicon cDNA sequence (5'-3')
WT
CAGCAGTAGGGGAACTTCTCCTGCTAGAATGGCTGGC
KR+TRS
CAGCTCTAAACGAACTTCTCCTGCTAGAATGGCTGGC
KR-TRS
CAGCAGTAAGAGGACTTCTCCTGCTAGAATGGCTGGC
M210I
CAGCAGTAGGGGAACTTCTCCTGCTAGAATTGCTGGC
Probe Sequence (5'-3) Target
WT: CAGCAGTAGGGGAAC Site 1
KR+TRS: CAGCTCTAAACGAAC Site 1
KR-TRS: CAGCAGTAAGAGGAC Site 1
WT-M210: TGCTAGAATGGCTGGC Site 2
Mut-M210I: TGCTAGAATTGCTGGC Site 2
D
Cq value
Probe
WT
KR+TRS
KR-TRS
WT-M210
Mut-M210I
NTC
Mock
rSARS2-WT
rSARS2-KR+TRS
rSARS2-KR-TRS
rSARS2-M210I
Samples
E
Proportion of population (%) [extracellular RNA]
hpi
WT
KR+TRS
**
F
ns
*
**
KR+TRS
KR-TRS
G
ns
WT
M210I

**Figure S2. rSARS-CoV-2 validation and extracellular RNA co-infection competition assay in HUVEC$^{ACE2}$.**

**A.** sgRNA profile of recombinant SARS-CoV-2 was determined by infecting HUVEC$^{ACE2}$ cells (MOI=4). RNA was harvested 24 hpi and subjected to RT-PCR and agarose gel electrophoresis.

**B.** N proteoform profile of SARS-CoV-2 variants was determined by infecting HUVEC$^{ACE2}$ cells (MOI=4). Protein lysate was harvested 24 hpi and subjected to SDS-PAGE and immunoblotting with anti-N.

**C.** Schematic of DNA probes used to differentiate rSARS-CoV-2 viruses. Probe and respective target sequence to either Site 1 (to differentiate WT, KR$^{+TRS}$, and KR$^{-TRS}$) or Site 2 (to differentiate WT from M210I).

**D.** TaqMan probe specificity was validated in RT-qPCR assays using cDNA as a template, generated from RNA from rSARS-CoV-2-infected HUVEC$^{ACE2}$ cells (24 hpi, MOI=4). All possible combinations of probes and recombinant virus-infected samples were tested alongside a no-template control (NTC); resulting $C_q$ values are displayed on a heatmap where reactions without amplification were set to $C_q = 40$.

**E-G.** Primary HUVEC$^{ACE2}$ cells were coinfected with equal infectious titers of the indicated recombinant virus to achieve a total MOI of 0.02. Time 0 represents the inferred proportions of each recombinant based on infectious titer input. At the indicated time post-infection, virus-containing supernatant was harvested, cell debris was removed by centrifugation (5 mins at 1000 RPMs), and extracellular RNA was harvested and subjected to probe-based RT-qPCR to differentiate recombinant virus abundance. These data represent three independent biological replicates (*n=3*). Statistics were performed using ratio paired T-test (*, $p < 0.0332$, ** $p < 0.0021$), standard error mean; SEM.

Figure S3

A

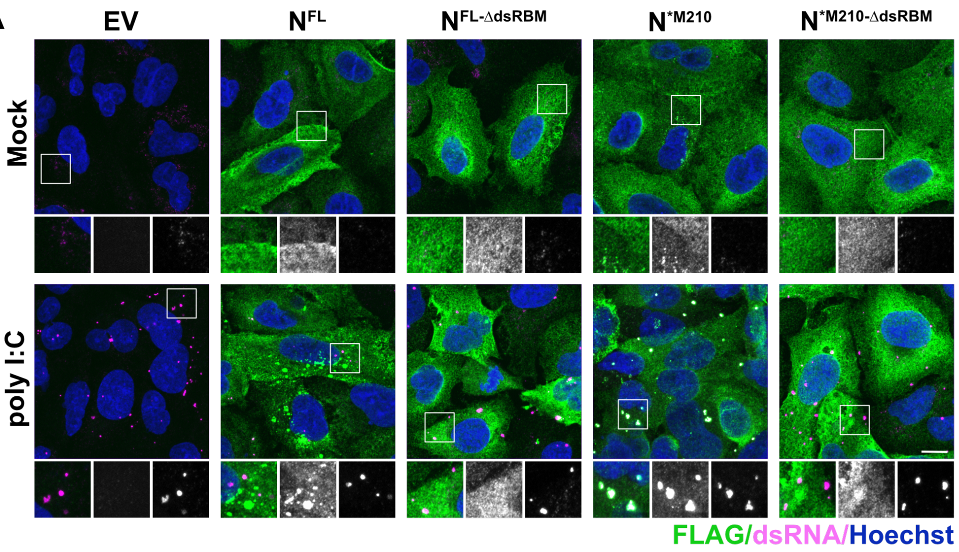
EV
N^FL
N^FL-ΔdsRBM
N^*M210
N^*M210-ΔdsRBM
Mock
poly I:C
FLAG/dsRNA/Hoechst

B

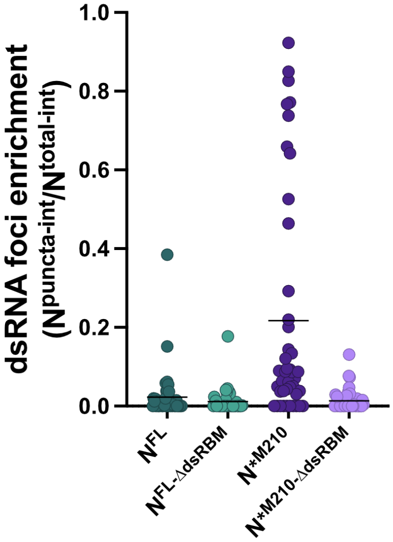
dsRNA foci enrichment
(N^puncta-int/N^total-int)
1.0
0.8
0.6
0.4
0.2
0.0
N^FL
N^FL-ΔdsRBM
N^*M210
N^*M210-ΔdsRBM

**Figure S3. $N^{*M210}$ is superior at sequestering dsRNA compared to $N^{FL}$.**

**A.** A549 cells were transduced with recombinant lentiviruses to ectopically express $N^{FL+/-dsRBM}$ or $N^{*M210+/-dsRBM}$ or EV. For all N proteoforms, internal methionine residues were mutated to ensure only the indicated proteoform of N was expressed ([$N^{FL}$; M210I and M234V], [$N^{*M210}$; M234V]). 96 hours post-transduction, cells were transfected with 0.5 µg high molecular weight poly I:C or mock transfected. Three hours post transfection, cells were fixed and immunostained with the FLAG antibody (N proteoform; Alexa 488) and J2 antibody (dsRNA; Alexa 647). Nuclei were stained with Hoechst. A maximum intensity projection is presented here. One representative experiment of two independent replicates is shown. Scale bar = 10 µm.

**B.** Enrichment of N proteoforms with RNA as in Fig 3B. These data represent two independent biological replicates (*n*=2) with 20 cells measured per condition, per replicate.

Figure S4

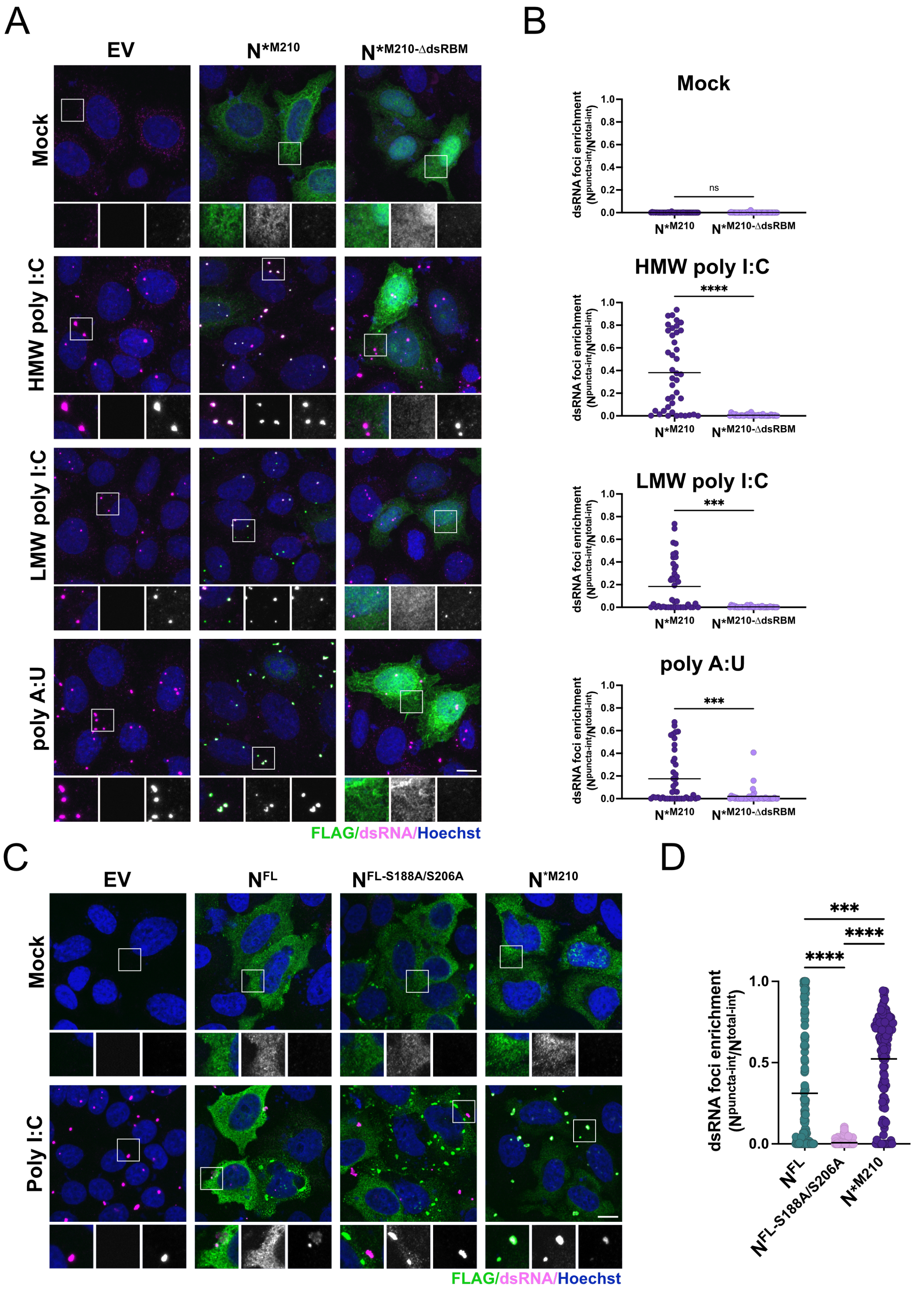
A
EV
N*M210
N*M210-ΔdsRBM
Mock
HMW poly I:C
LMW poly I:C
poly A:U
FLAG/dsRNA/Hoechst
B
Mock
HMW poly I:C
LMW poly I:C
poly A:U
dsRNA foci enrichment (Npuncta-int/Ntotal-int)
ns
****
***
***
N*M210
N*M210-ΔdsRBM
C
EV
NFL
NFL-S188A/S206A
N*M210
Mock
Poly I:C
FLAG/dsRNA/Hoechst
D
dsRNA foci enrichment (Npuncta-int/Ntotal-int)
***
****
****
NFL
NFL-S188A/S206A
N*M210

**Figure S4. Hypo-phosphorylation of $N^{FL}$ reduces dsRNA interaction.**

**A.** EV, $N^{*M210}$, or $N^{*M210\text{-}\Delta dsRBM}$-expressing HeLa cells were transfected with 0.5 µg of high molecular weight (HMW) or low molecular weight (LMW) poly I:C or poly A:U, or mock transfected. Three hours post transfection, cells were fixed and immunostained with the FLAG antibody (N proteoform; Alexa 488) and J2 antibody (dsRNA; Alexa 647). Nuclei were stained with Hoechst. A maximum intensity projection (MIP) is presented here. One representative experiment of three independent replicates is shown ($n$=3). Scale bar = 10 µm.

**B.** Enrichment of N proteoforms with dsRNA as in Fig 3B. These data represent three independent biological replicates ($n$=3) with 18 cells measured per condition, per replicate. Statistics were performed using a Mann-Whitney test (***, $p < 0.0002$, ****, $p < 0.0001$).

**C.** EV, $N^{FL}$, $N^{FL\text{-}S188A/S206A}$, or $N^{*M210}$-expressing HeLa cells were transfected with 0.5 µg poly I:C or mock transfected. Three hours post transfection, cells were fixed and immunostained with the FLAG antibody (N proteoform; Alexa 488) and J2 antibody (dsRNA; Alexa 647). Nuclei were stained with Hoechst. A maximum intensity projection (MIP) is presented here. One representative experiment of three independent replicates is shown ($n$=3). Scale bar = 10 µm.

**D.** Enrichment of N proteoforms with dsRNA as in B. These data represent three independent biological replicates ($n$=3) with 40 cells measured per condition, per replicate. Statistics were performed using a Kruskal-Wallis $H$ test with Dunn's correction (****, $p < 0.0001$), mean.

Figure S5

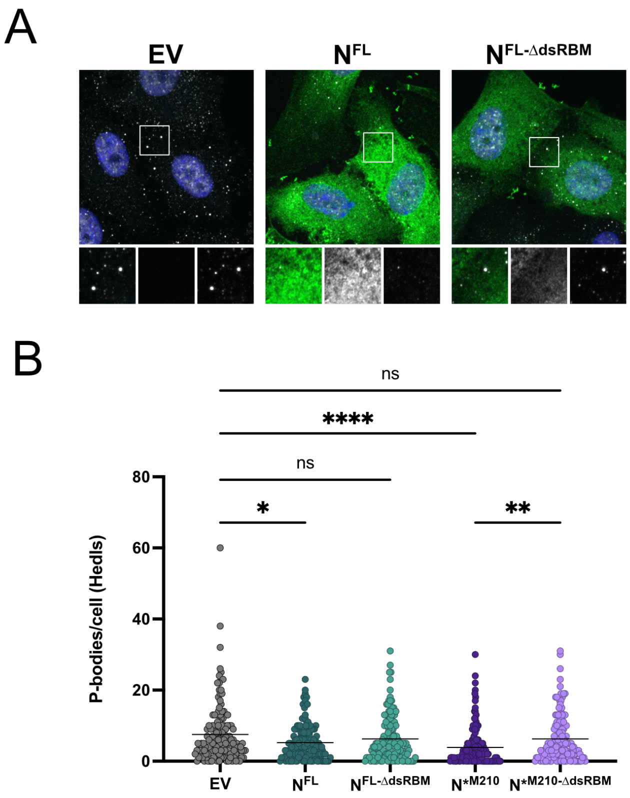
A
EV
N^FL
N^FL-ΔdsRBM
N^*M210
N^*M210-ΔdsRBM
FLAG/Hedls/Hoechst
B
ns
****
ns
*
**
80
60
40
20
0
P-bodies/cell (Hedls)
EV
N^FL
N^FL-ΔdsRBM
N^*M210
N^*M210-ΔdsRBM

**Figure S5. The dsRBM of $N^{*M210}$ is required for P-body disassembly.**

**A.** Primary HUVEC cells were transduced with recombinant lentiviruses to ectopically express $N^{FL+/-dsRBM}$, $N^{*M210+/-dsRBM}$, with internal methionines mutated ([$N^{FL}$; M210I and M234V], [$N^{*M210}$; M234V]), or EV. 96 hours post-transduction, cells were fixed and immunostained with the FLAG antibody (N proteoform; Alexa 488) and the Hedls antibody (P-bodies; Alexa 647). Nuclei were stained with Hoechst. A maximum intensity projection (MIP) is presented here. One representative experiment of three independent replicates is shown. Scale bar = 10 µm.

**B.** P-bodies were quantified using CellProfiler by measuring Hedls puncta in N-expressing cells (thresholded by FLAG staining) or EV transduced cells. These data represent three independent biological replicates ($n$=3) with >20 cells measured per condition, per replicate. Each datapoint represents a single cell. Statistics were performed using a Kruskal-Wallis $H$ test with Dunn's correction (*, $p < 0.032$, **, $p < 0.0021$, ****, $p < 0.0001$).
